# The impact of sex and physical activity on the local immune response to muscle pain

**DOI:** 10.1101/2022.12.07.519473

**Authors:** Joseph B. Lesnak, Kazuhiro Hayashi, Ashley N. Plumb, Adam J. Janowski, Michael S. Chimenti, Kathleen A. Sluka

## Abstract

Induction of muscle pain triggers a local immune response to produce pain and this mechanism may be sex and activity level dependent. The purpose of this study was to measure the immune system response in the muscle following induction of pain in sedentary and physically active mice. Muscle pain was produced via an activity-induced pain model using acidic saline combined with fatiguing muscle contractions. Prior to induction of muscle pain, mice (C57/BL6) were sedentary or physically active (24hr access to running wheel) for 8 weeks. The ipsilateral gastrocnemius was harvested 24hr after induction of muscle pain for RNA sequencing or flow cytometry. RNA sequencing revealed activation of several immune pathways in both sexes after induction of muscle pain, and these pathways were attenuated in physically active females. Uniquely in females, the antigen processing and presentation pathway with MHC II signaling was activated after induction of muscle pain; activation of this pathway was blocked by physical activity. Blockade of MHC II attenuated development of muscle hyperalgesia exclusively in females. Induction of muscle pain increased the number of macrophages and T-cells in the muscle in both sexes, measured by flow cytometry. In both sexes, the phenotype of macrophages shifted toward a pro-inflammatory state after induction of muscle pain in sedentary mice (M1+M1/2) but toward an anti-inflammatory state in physically active mice (M2+M0). Thus, induction of muscle pain activates the immune system with sex-specific differences in the transcriptome while physical activity attenuates immune response in females and alters macrophage phenotype in both sexes.

## Introduction

Chronic pain is a major health care issue in the United States affecting up to 100 million Americans (*1*). Chronic musculoskeletal pain is the most common type of chronic pain and results in significant disability and loss of function (*1–4*). Women report higher prevalence and severity of symptoms (*3, 4*) yet, the underlying mechanisms for the development of musculoskeletal pain have primarily been investigated in males (*5*). Recent evidence in animals with neuropathic, prolactin induced, and incisional pain models have uncovered sex specific mechanisms particularly in the immune system (*5–8*). It has been suggested males develop pain through activation of the innate immune system (i.e. microglia, macrophages) while females develop pain through the adaptive immune system (i.e. T-cells) (*9*). However, there is conflicting evidence as prior research has supported a role of T-cells for the generation of pain in males, which might depend on pain model, location studied, and strain of animal utilized (*10*).

Our prior work shows muscle macrophages play a key role in the generation of muscle pain and the ability of physical activity to prevent it. Depletion of muscle macrophages prior to induction of the pain model prevents development of muscle hyperalgesia (*11–13*), and depletion of P2X4 from muscle macrophages prevents hyperalgesia in both male and female mice (*14*). Similarly, in models of neuropathic pain, macrophages at the site of injury play a key role in the production of hyperalgesia (*15, 16*), suggesting local immune mechanisms are important for pain development. Further, regular physical activity alters macrophage phenotype in skeletal muscle by increasing the number of anti-inflammatory macrophages (M2), and blockade of antiinflammatory cytokine receptors (IL-4, IL-10) at the site of insult prevents the analgesia produced by regular physical activity (*13, 17–20*); however, these studies were only done in male mice. These data suggest macrophages at the site of insult play a dual role in the generation and prevention of pain. A more complete understanding of the local immune system response, and potential differences between the sexes, could lead to the development of novel therapeutic targets.

Next generation RNA sequencing analyzes differences in the transcriptome and allows for the discovery of novel pain mechanisms in an unbiased manner. RNA sequencing has been utilized on mouse skeletal muscle to identify activation of genes and pathways in models of injury and age-related degeneration, but not in the context of muscle pain (*21–23*). Recently, sex-specific transcriptional alterations have been uncovered utilizing RNA sequencing following induction of models of neuropathic and prolactin induced pain that highlight the need to explore for sex differences in pain research (*7, 8*). Therefore, the goal of this study was to use RNA sequencing and flow cytometry to explore sex-specific alterations in the immune system following induction of muscle pain in sedentary and physically active animals. We identified a female specific upregulation of genes associated with immune system function of antigen processing and presentation induced by muscle pain that was attenuated by regular physical activity, suggesting a role for the adaptive immune system in production of muscle pain.

## Methods

### Study and Experimental Design

This study was designed as a randomized, controlled laboratory experiment using C57BL/6J mice. In each experiment, equal numbers of male and female mice were used. All animals were randomly allocated to experimental groups using a random number generator in blocks of 4, stratified by sex. To control for potential batch effects in RNA sequencing experiments, equal numbers of animals from each sex and experimental group were collected within each day of tissue harvesting. The main goal of this study was to uncover sex specific immune system mechanisms in the production of muscle pain and in physical activity’s ability to prevent muscle pain through the use of RNA sequencing and flow cytometry.

RNA sequencing was performed on mouse skeletal muscle following induction of a muscle pain model or from pain free controls. Mice were first allocated as sedentary or physically active. Sedentary mice were further allocated to receive the activity-induced muscle pain model (n=8) or be pain free (n=8), while all physically active animals received the muscle pain model (n=8). The left gastrocnemius muscle was harvested 24 hours after induction of the pain model and RNA sequencing was performed (Fig. 1A). The primary endpoint for this experiment was group and sex differences in differentially expressed genes (DEG’s).

**Figure 1.**
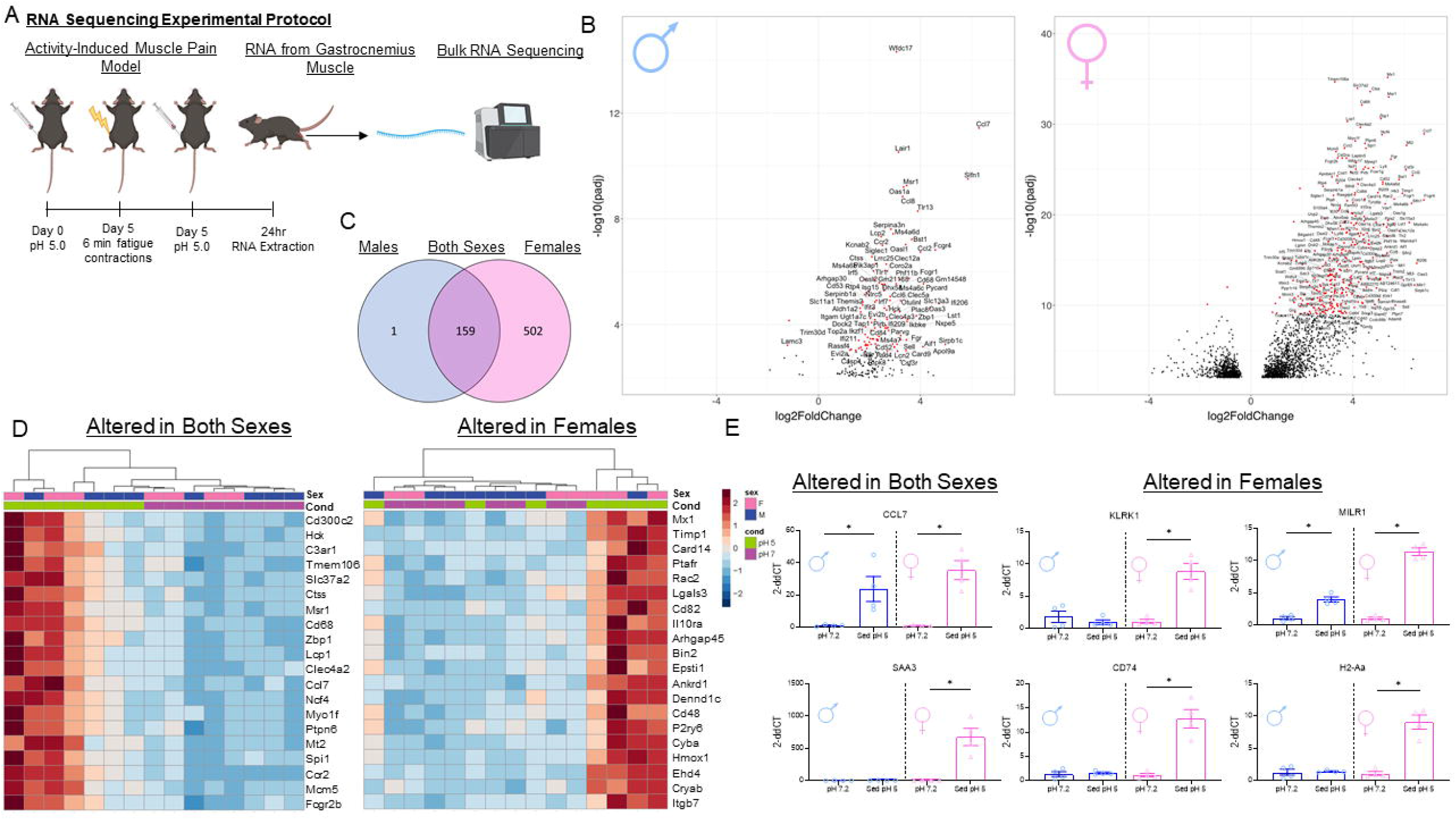
RNA sequencing reveals sex differences in the number of DEGs in the gastrocnemius muscle between pain and pain free animals. A. Graphical depiction of experimental protocol of RNA sequencing experiment. B. Volcano plots of differentially expressed genes (DEGs) between animals who received the activity-induced muscle pain model and pain free animals for males (left) and females (right). Only genes with an adjusted p-value <0.1 are plotted with red dots depicting a more significant p-value (Males: adj-p<0.001; Females: adj-p<1e-9). Female mice had a higher average absolute magnitude of fold change and number of DEG’s compared with males. C. When comparing between the sexes, 159 genes were differentially expressed in both sexes, while 1 gene in males and 502 genes in females were differentially expressed in that sex only. D. Heat maps for top 20 DEG’s with the smallest adjusted p-values for genes altered in both sexes (left) and genes altered in females only (right). D. qPCR confirmation of select genes from RNA sequencing analysis. For DEG’s from both sexes *CCL7* but not *SAA3* was found to be upregulated in both male and female mice. For DEG’s in females only, *KLRK1, CD74*, and *H2-Aa* were confirmed to be increased in females only, while *MILR1* was found increased in both sexes. *p<0.05 compared with pH 7.2; Data are mean+SEM; Images made on BioRender.

Results of RNA sequencing were first validated with qPCR. Again, mice were allocated as pain free or sedentary or physically active with the pain model (n=8 per group). The left gastrocnemius muscle was harvested 24 hours after induction of the pain model. A total of 10 genes, 6 for pain free versus sedentary with the pain model and 4 for sedentary versus physically active animals, were selected from the RNA sequencing results and expression levels were measured via qPCR.

Based on female specific increased signaling of MHC II from RNA sequencing results, we tested if blockade of MHC II would prevent the development of muscle pain in male and female mice. Muscle hyperalgesia was assessed at baseline and 24 hours after induction of the muscle pain model. To block MHC II, mice were allocated to receive either an MHC II antibody (n=16) or its control (n=16) 30 minutes prior to fatiguing muscle contractions.

We utilized flow cytometry to further validate RNA sequencing results and explore changes in immune cell populations following induction of muscle pain. Mice were allocated to pain free or sedentary or physically active with the pain model (n=8 per group) and the left gastrocnemius muscle was harvested 24 hours after induction of the pain model and flow cytometry was performed the same day.

Lastly, we tested whether modulation of testosterone altered the expression of female specific DEG’s from RNA sequencing analysis. Testosterone levels were altered in male and female mice 2 weeks prior to induction of the activity-induced muscle pain model. In males, animals received an orchiectomy (n=6), sham orchiectomy (n=6), or were pain free without a surgery (n=6). In females, animals received implantation of a slow-release testosterone (7.5mg) (n=6), a vehicle pellet (n=6) or were pain free without a surgery (n=6). Left gastrocnemius muscle was harvested 24 hours after induction of the pain model. Four female specific DEG’s were selected from the RNA sequencing results and expression levels were measured via qPCR.

### Animals

A total of 140 C57BL/6J mice (70 male, 70 female) (20-30g) (8 weeks of age) (Jackson Laboratories, Bar Harbor, ME, USA) were used in the following experiments. All mice were housed on a 12-hour light-dark cycle with access to food and water *ad libitum*. All animal protocols were approved by the University of Iowa Animal Care and Use Committee and were conducted in agreement with National Institute of Health guidelines.

### Activity-Induced Muscle Pain Model

The activity-induced muscle pain model was produced as previously described (*24*). Briefly, this pain model uses a combination of intramuscular (i.m.) acidic saline injections and fatiguing muscle contractions. On day zero, mice were anesthetized with 2-4% isoflurane via vaporizer and given an i.m. injection of 20μl of pH 5.0±0.1 saline into the left gastrocnemius muscle. On day five, mice were anesthetized with 2-4% isoflurane via vaporizer and needle electrodes were inserted into the left gastrocnemius muscle in parallel orientation. To induce fatigue, 6 minutes of submaximal contractions were delivered at 40 Hz with an amplitude of 7V and a duty cycle of 3.75 seconds on and 4.25 seconds off. The fatiguing muscle contractions produces an approximate 60% reduction in muscle force and a significant decrease in muscle pH (*24*). This pain model produces a sex-dependent pain phenotype in which males develop unilateral muscle hyperalgesia and females develop bilateral muscle hyperalgesia (*24–26*). Pain free controls were produced by delivering an i.m. injection of 20μl of pH 7.2±0.1 saline into the left gastrocnemius muscle on days zero and five.

### Physical Activity

Physical activity was increased in animals using voluntary wheel running. Animals in the physically active group were individually housed with a running wheel in their home cage with 24 hour access for 8 weeks. Voluntary wheel running allows animals to dictate the timing, speed, and duration of running and avoids increased stress demonstrated in forced exercise models (*27*). To mimic housing conditions of physically active mice, sedentary and pain free animals were individually housed for 8 weeks. Following 8 weeks, the activity-induced muscle pain model was applied. The running wheels were removed from cages at the beginning of the pain model induction (first pH 5.0 injection). We have repeatedly shown 8 weeks of voluntary wheel running prior to induction of a muscle pain prevents the onset of muscle hyperalgesia in male and female mice (*17, 18, 20, 28*). Total number of wheel revolutions was recorded over the 8 weeks of physical activity (Supplemental Table 1).

### RNA Extraction

Animals were euthanized with CO_2_ and the left gastrocnemius muscle was harvested and placed in RNAlater (Invitrogen, Thermofisher Scientific, Waltham, MA, USA). RNA was isolated using a fibrous tissue mini kit (Qiagen, Hilden, Germany) and further purified with an RNA clean and concentrate kit (Zymo Research, Orange, CA, USA) according to the manufacturer’s instructions. Only RNA samples with A260/A280 values greater than 1.8 and RNA integrity numbers greater than 8.0 were used for RNA sequencing or qPCR analysis. RNA was stored at −80°C prior to subsequent analysis.

### RNA Sequencing

Barcoded samples were pooled and sequenced, using an Illumina NovaSeq 6000 at the Iowa Institute for Human Genomics (IIHG) Core Facility, to obtain a minimum of 30 million paired-end 150bp reads per sample, as per ENCODE standards (*29, 30*). Reads were converted from the native Illumina BCL format to fastq format with ‘*bcl2fastq*’ (v2.20.0.422). The ‘*bcbio-nextgen*’ python pipeline (https://github.com/chapmanb/bcbio-nextgen; v.1.2.4) was run in “RNA-seq” mode to align reads against the Ensembl mm10/GRCm38.86 reference build using the splice-aware ‘*hisat2*’ aligner (v.2.2.1) (*31, 32*). Alignments were stored as indexed BAM files. Quantification of reads against the mouse transcriptome (GENCODE M25) was also performed using the *salmon* aligner (*33*). Analysis of BAM alignments showed that for all samples, >90% of reads were mapped as proper pairs to the reference, with >93% of mapped reads originating from exonic regions. Additional QC was performed with *MultiQC (34)*. All samples passed bioinformatic QC thresholds and were retained for exploratory analysis. Salmon-derived transcript expression values were summarized to the gene level using *tximport* (*35*). Normalized and variance-stabilized gene-level counts were used for exploratory analysis via principal components analysis (PCA) and sample-to-sample distance calculations. One sample, so-called “GM-45” (female; sedentary, activity-induced pH 5 pain model) showed extreme outlier behavior in the PCA and sample distance analysis (Supplemental Fig. 1+2). The outlier was confirmed with poor laboratory RNA quality scores (data not shown) and the sample was dropped from the downstream analysis. Unnormalized *salmon* gene-level counts were used for differential expression analysis with *DESeq2* (*36*) following best practices as described in the DESeq2 vignette (*37*). Samples were split on animal sex and separate male and female DESeq2 data objects were created and conditioned on tissue batch and experimental condition for DE testing. Adjusted p-values were calculated using a Benjamini, Hochberg correction with the threshold for acceptance set at 10% (*38*). Code used in this analysis is available in an online repository (https://github.com/mchimenti/project_rnaseq_mouse_21_056_Jun2021/blob/main/analysis_21_056_sampleset_SplitSex_eda_deseq2_jun2021.Rmd). Raw and processed data were deposited in the NCBI Gene Expression Omnibus (GSE206187). Pathway analysis was carried out using iPathwayGuide software (https://advaitabio.com/ipathwayguide) to perform proprietary “impact analysis” to generate reports describing enriched pathways, Gene Ontology (GO) terms, and upstream regulatory genes. Pathway analysis examines patterns of DEGs that are found to be enriched in specific biological pathways, Gene Ontology divides the function of genes into biological processes, molecular function, and cellular components, and upstream regulators are genes that are predicted as being present due to their differentially expressed target genes. These analyses consider both the direction and type of all signals on a pathway along with the position, role, and type of each gene (*39–41*). A correction for False Discovery Rate was used to analyze iPathwayGuide data.

### qPCR

After gastrocnemius muscles were harvested and RNA was extracted, reverse transcription was carried out with 500ng of RNA using an Affinity Script qPCR cDNA Synthesis Kit (Agilent Technologies, Santa Clara, CA, USA) according to manufacturer’s instructions. The cDNA was stored at −20°C until qPCR analysis was performed. cDNA was used as a template for qPCR using Power SYBR Green PCR Master Mix (Applied Biosystems, Thermo Fisher Scientific, Waltham, MA, USA). All forward and reverse primers were developed using NCBI Primer-BLAST software to ensure no off-target amplification (http://www.ncbi.nlm.nih.gov/tools/primer-blast/) and are listed in Supplemental Table 2. All primers were manufactured by Integrated DNA Technologies (Coralville, IA, USA). Analysis of qPCR amplifications was performed with QuantStudio 7 Flex Real-Time PCR System (Applied Biosystems) using the 2-ΔΔCT method with 36B4 serving as an internal control. All samples were run in duplicate, and the cycle threshold was averaged between the two samples. Reactions were performed under the following conditions: 2 min at 50°C, 10 min at 95°C, 40 cycles of 15 sec at 95°C and 1 min at 60°C.

### Behavioral assessment

Muscle hyperalgesia was assessed by determining muscle withdrawal thresholds (MWT). Mice were restrained in a gardener’s glove and a custom built, force sensitive tweezer was used to apply a force to the left and right gastrocnemius muscle as previously described (*42, 43*). The gastrocnemius muscle was squeezed until the animal withdrew its hindlimb or made an auditory response. Both the left and the right gastrocnemius muscle were tested and an average of 3 trials were used to determine MWT for each limb. The presence of muscle hyperalgesia is indicated by a decrease in MWT.

### Blockade of MHC II

Blockade of MHC II during induction of the activity-induced muscle pain model was performed through the utilization of an anti-mouse MHC II in vivo antibody. First, animals were anesthetized with 2-4% isoflurane via vaporizer. Then animals received a 20μl i.m. injection of either MHC II antibody (BE0108, Bio X Cell, Lebanon, NH, USA) or an IgG2b isotype control (BE0090, Bio X Cell) at a concentration of 10mg/ml into the left gastrocnemius muscle. Fatiguing muscle contractions of the left gastrocnemius muscle were initiated as described above 30 minutes following MHC II antibody or IgG control injections.

### Flow Cytometry

Animals were euthanized with CO_2_ and the left gastrocnemius muscle was harvested and placed in 0.2% Collagenase Type II (Worthington, Lakewood, NJ, USA) dissolved in Dulbecco’s Modified Eagle Medium (DMEM; Gibco, Waltham, MA, USA). Muscle tissue was then diced with small scissors and incubated at 37°C for 20 minutes. Muscle samples were then triturated with a 5ml stripette and incubated for another 20 minutes at 37°C. Ice cold wash buffer (50% volume Hanks Balanced Salt Solution (HBSS; Gibco), 50% volume Accumax (Accutase, San Diego, CA, USA), 0.1mg/ml DNase (Roche, Millipore Sigma, St. Louis, MO, USA)) was added to each sample and triturated with a 10ml stripette. Samples were then filtered through 70μm nylon mesh strainer. Samples underwent two wash cycles by being centrifuged at 300 × g at 4°C for 5 minutes with resuspension in wash buffer. After the second wash, samples were resuspended in flow cytometry staining buffer (eBioscience, Invitrogen, Waltham, MA, USA). Samples were incubated on ice with an Fc receptor blocker (TruStain FcX PLUS (anti-mouse CD16/32) Antibody)) for 10 minutes. Samples were then incubated with conjugated flow cytometry antibodies (Supplemental Table 3) on ice for 20 minutes in the dark. Samples then underwent two wash steps by being centrifuged at 300 x g at 4°C for 5 minutes with resuspension in flow cytometry staining buffer. After the second wash, samples were resuspended in flow cytometry staining buffer with DNase (Roche; 1mg/ml) and left on ice, in the dark, until they were analyzed. Appropriate reference controls were used for unmixing within each run of flow cytometry data collection. Fluorescence Minus One (FMO) controls were used to determine optimal gates. Hoechst was used to determine intact cells and then further gated with CD45 to determine immune cells. To determine immune cell populations, cells were gated for CD3 (CD3+), Ly6C (CD3-,Ly6C+), CD205 (CD3-, Ly6C-,CD205+), and CD49b (CD3-,Ly6C-, CD4bb+). CD3+ cells were subsequently gated for CD4, and CD8. Ly6C+ cells were subsequently gated for F4/80, CD206, and CD11c. An MHC II antibody was used to explore differences in MHC II on macrophages and immune cells. Flow cytometry was performed on a Cytek Aurora five-laser spectral cytometer (16-UV, 16-Violet, 14-Blue, 10-Yellow/Green and 8-Red laser configuration) (CytekBio, Freemont, CA, USA) and data was analyzed with FlowJo v10.8.1(Becton, Dickinson & Company, Franklin Lakes, NJ, USA).

### Orchiectomy

Male animals were anesthetized with 2-4% isoflurane via vaporizer. An incision was made over the scrotum, the testes were exposed, and the vas deferens and testicular blood vessels were ligated with sutures. The testes were then removed, and the scrotal incision was closed with sutures. Sham orchiectomy surgery was performed by exposing the testes through a scrotal incision but not removing them. After surgery, all animals were moved to individual cages to protect the surgical site.

### Testosterone Administration

Female animals were anesthetized with 2-4% isoflurane via vaporizer. An incision was made over the nape of the neck and a slow-release testosterone (7.5mg, 60-day release; Innovative Research of America, Sarasota, FL, USA) or vehicle pellet was implanted subcutaneously. The incision was closed with sutures. After surgery, all animals were moved to individual cages to protect the surgical site.

### Statistical Analysis

All data is presented as mean±SEM. For qPCR data a t-test was used to compare differences between groups. To analyze behavior data, a generalized linear model with fixed effects for group and sex was utilized for the 24 hour MWT value using baseline as a covariate. If a significant interaction for group and sex was found, male and female data were separated, and a t-test was utilized to explore group differences for the 24 hour MWT value. A Cohen’s d effect size was calculated for 24 hour MWT values. For behavior data we also calculated a MWT value change score by subtracting the 24 hour value from the baseline value. A t-test was utilized to determine group differences for MWT change score values. Male contralateral limb values were not analyzed since in males, the activity-induced muscle pain model fails to produce muscle hyperalgesia on the contralateral limb (*24*). For flow cytometry data, a generalized linear model with fixed effects for group and sex was used, with a Tukey post hoc test applied to further determine group differences. If a significant interaction for group and sex was found, male and female data were separated, and a one-way ANOVA was utilized to explore for group differences followed by a Tukey post hoc test when appropriate. For RNA sequencing analysis, a gene needed to have an adjusted p-value of <0.1 and an absolute log2 fold change of >2.0 to be considered a DEG. For all other experiments, a p-value of <0.05 was considered statistically significant. Statistical analyses were performed on SPSS Version 27.0 (SPPS Inc. Chicago, IL, USA) and GraphPad Prism Version 9.00 (GraphPad Software, La Jolla, CA, USA).

## Results

### Sex differences in muscle transcriptome following induction of activity-induced muscle pain

To determine if there are sex differences in the muscle transcriptome following induction of muscle pain, we used bulk RNA sequencing of mouse skeletal muscle 24 hours after induction of the model (Fig. 1A). To determine sex differences, differential expression analysis was performed by examining the male and female data separately and then comparing results between the sexes.

The total number of genes with measured expression was 13,621 in both males and females. Principal component analysis (PCA) demonstrated that one female sample from the sedentary in pain group displayed outlier behavior and was subsequently dropped from the RNA sequencing analysis leaving a sample size of 3 for this group. The PCA plots demonstrate that samples clustered together based on treatment group and sex which was expected based on our hypothesis of sex differences in pain mechanisms (Supplemental Fig. 1A+B). The PCA plots also display lack of clustering based on date of tissue collection suggesting the absence of any batch effect in the RNA sequencing data (Supplemental Fig. 1C).

In males we identified 160 genes with differential expression between sedentary in pain and pain free animals of which all 160 were upregulated (adjusted p-value <0.1; abs(log_2_FC)>2) (Fig. 1B Left Panel). In females, 661 genes were identified with differential expression of which 591 were upregulated and 70 were downregulated between sedentary in pain and pain free animals (adjusted p-value <0.1; abs(log_2_FC>2)) (Fig. 1B Right Panel). Females also had a greater average absolute log2 fold change of DEGs when compared with males (Female: 3.22±0.98, Male: 2.77±0.88). Of the DEGs, 159 were differentially expressed in both sexes, 1 was altered in males only (*Atp1a3*), and 502 were altered in only females (Fig. 1C). Heat maps for the top DEG’s based on adjusted p-value for both sexes and females are displayed in Figure 1D. Heat maps for DEG’s in both sexes demonstrate a clear separation of sedentary in pain (pH 5) and pain free (pH 7) groups (Fig. 1D Left Panel) while female only DEG’s demonstrate a separation between males and females within each treatment group (Fig. 1D Right Panel).

Overall, females saw a greater increase in the number and magnitude of fold change of DEGs following induction of the activity-induced muscle pain model when compared with males.

To validate RNA sequencing results, 6 genes were selected and measured with qPCR in a separate set of animals within the same experimental groups. These genes were chosen based on their high magnitude of fold change and their prevalence in subsequent pathway analysis. Two genes that were upregulated in males and females (*CCL7, SAA3*) and four genes that were upregulated in females only (*KLRK1, MILR1, CD74, H2-Aa*) were tested. For the genes upregulated in both sexes from our RNA sequencing results, *CCL7* was increased in males and females following induction of the activity-induced muscle pain model (Male: p=0.02; Female: p<0.01), while *SAA3* was increased in females only (Male: p=0.99; Female: p<0.01) (Fig 1E). For the genes upregulated in females only from our RNA sequencing results, *KLRK1, CD74*, and *H2-Aa* were increased in females but not males (Male: p=0.87-0.99; Female: p<0.01) while *MILR1* was increased in both sexes (Male: p<0.01; Female: p<0.01) following induction of activity-induced muscle pain (Fig. 1E). Thus, we were able to validate RNA sequencing results via qPCR in a separate set of animals in 10 out of the 12 comparisons across the 6 tested genes.

### Pathway Analysis Reveals Sex Differences in Immune System Activation

To further investigate the potential function of the specific DEGs in this data set, pathway analysis, Gene Ontology (GO) terms, and upstream regulators were explored using iPathwayGuide software (AdvaitaBio). For pathway analysis we report the most significant enriched pathways based on adjusted p-value regardless of sex. Since males had only one unique gene upregulated, we report the most significant GO terms and upstream regulators based on adjusted p-value that were found to be significant in both sexes and in females only.

Overall, the majority of the pathways, GO terms, and upstream regulators that increased in both males and females involve immune system function including the innate immune response, chemokine signaling, and toll-like receptor signaling (Fig. 2A) (Tables 1–3). Of note, we found a robust sex difference in the antigen processing and presentation pathway which encompasses major histocompatibility 1 and 2 signaling (MHC I and MHC II). Females demonstrated activation of 17/50 genes while males demonstrated upregulation in 2/50 genes in the antigen processing and presentation pathway (Fig. 2A-C). Females, but not males, had a significant upregulation of several genes associated with MHC II signaling including *CD74* and *H2-Aa* which we confirmed with qPCR to be upregulated in females only (Fig. 1E). Confirmation of upregulation of the MHC II signaling pathway in females was also demonstrated by increased GO terms involving MHC II in female but not male muscle (Bolded and italicized in Tables 1–3 and Fig 2A). Lastly, upstream regulator analysis demonstrated DEGs were consistent with signaling of cytokines IL-1β and IL-33 in both sexes and with INFγ, C-reactive protein, and colony stimulating factor in females (Fig. 2D). Females showed 18 DEGs consistent with the presence of INFγ while males had 3. This also supports the female specific activation of antigen processing and presentation as INFγ has been demonstrated to increase MHC II on antigen presenting cells (*44*). In sum, this data suggests that males and females had upregulation of several immune system pathways in the muscle following induction of pain. However, females, but not males, had activation of the antigen processing and presentation pathway as well as MHC II signaling after induction of muscle pain.

**Figure 2.**
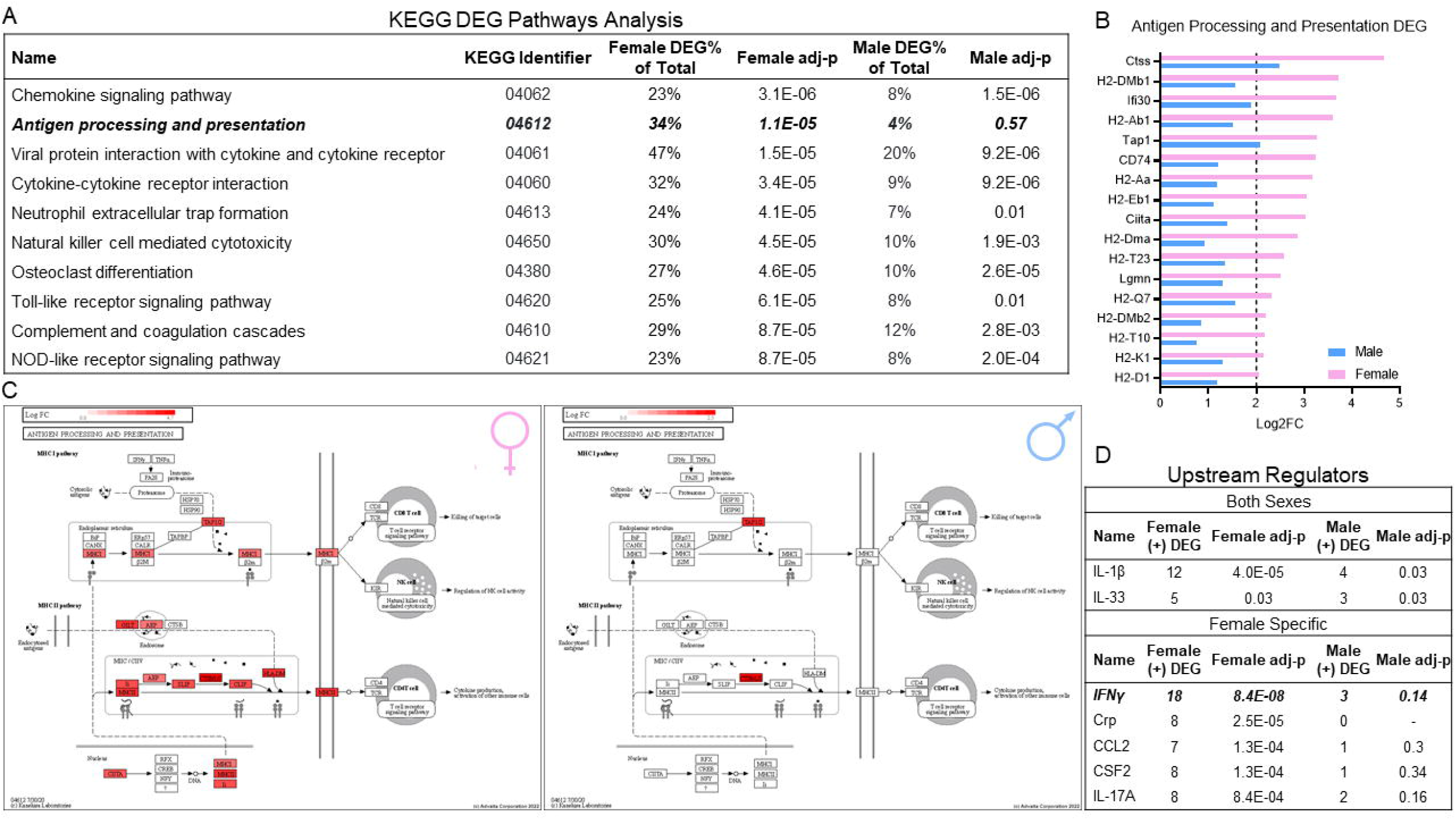
Pathway analysis reveals induction of muscle pain enriches several immune system pathways in both sexes but antigen processing and presentation in females only. A. Top 10 enriched pathways with smallest adjusted p-values from RNA sequencing analysis for both male and female sedentary mice who received the activity-induced muscle pain model. Antigen processing and presentation pathway was the only pathway in the top 10 that was sexually dimorphic with females, but not males, showing significant activation. B. Log2 fold change of the 17 genes upregulated in the antigen processing and presentation pathway for male and female mice. Only 2 of the 17 genes (*Ctss, Tap1*) were upregulated in male mice. C. Graphical depiction of antigen processing and presentation pathway for female (left) and males (right). Red represents differentially expressed genes throughout the pathway. D. Top upstream regulators with smallest adjusted p-value predicted to be present for both sexes and females only based on DEGs. IL=interleukin; IFNγ=interferon gamma; Crp= C reactive protein; CSF= colony stimulating factor

**Table 1.** Top 20 Gene Ontology biological processes based on adjusted p-value for both sexes and females only

Biological Processes
| Both Sexes |  |  |  |  |  | Female Specific |  |  |  |  |  |
| --- | --- | --- | --- | --- | --- | --- | --- | --- | --- | --- | --- |
| Name | GO Identifier | Female %DEG of Total | Female adj-p | Male %DEG of Total | Male adj-p | Name | GO Identifier | Female %DEG of Total | Female adj-p | Male %DEG of Total | Male adj-p |
| Innate immune response | GO:0045087 | 33% | 1.0E-24 | 11% | 1.0E-24 | Interleukin-1 beta production | GO:0032611 | 39% | 5.4E-16 | 4% | 0.15 |
| Defense response to other organism | GO:0098542 | 31% | 1.0E-24 | 10% | 1.0E-24 | Regulation of interleukin-1 beta production | GO:0032651 | 39% | 1.7E-15 | 5% | 0.13 |
| Regulation of immune response | GO:0050776 | 26% | 1.0E-24 | 8% | 1.0E-24 | Interleukin-1 production | GO:0032612 | 34% | 8.0E-15 | 4% | 0.19 |
| Response to external biotic stimulus | GO:0043207 | 26% | 1.0E-24 | 8% | 1.0E-24 | Production of molecular mediator of immune response | GO:0002440 | 23% | 2.6E-14 | 3% | 0.08 |
| Response to other organism | GO:0051707 | 26% | 1.0E-24 | 8% | 1.0E-24 | Regulation of interleukin-1 production | GO:0032652 | 34% | 2.7E-14 | 4% | 0.18 |
| Immune response | GO:0050776 | 25% | 1.0E-24 | 8% | 1.0E-24 | Positive regulation of interleukin-1 beta production | GO:0032731 | 45% | 2.3E-13 | 7% | 0.07 |
| Response to biotic stimulus | GO:0009607 | 25% | 1.0E-24 | 8% | 1.0E-24 | Response to protozoan | GO:0001562 | 60% | 2.4E-12 | 4% | 0.43 |
| Defense response | GO:0006952 | 26% | 1.0E-24 | 8% | 1.0E-24 | *Antigen processing and presentation of exogenous peptide antigen via MHC class II* | GO:0019886 | 80% | 3.6E-12 | 13% | 0.06 |
| Biological process involved in interspecies interaction between organisms | GO:0044419 | 24% | 1.0E-24 | 7% | 1.0E-24 | Positive regulation of interleukin-1 production | GO:0032732 | 40% | 3.8E-12 | 6% | 0.08 |
| Regulation of immune system process | GO:0002682 | 21% | 1.0E-24 | 7% | 1.0E-24 | Response to lipid | GO:0033993 | 13% | 2.0E-11 | 2% | 0.13 |
| Response to external stimulus | GO:0009605 | 18% | 1.0E-24 | 6% | 1.0E-24 | Positive regulation of adaptive immune response based on somatic recombination of immune receptors built from immunoglobulin superfamily domains | GO:0002824 | 28% | 3.4E-11 | 4% | 0.08 |
| Regulation of immune system process | GO:0002682 | 21% | 1.0E-24 | 7% | 1.0E-24 | Defense response to protozoan | GO:0042832 | 57% | 2.6E-10 | 4% | 0.40 |
| Response to interferon-beta | GO:0035456 | 69% | 1.0E-24 | 15% | 2.6E-05 | Negative regulation of cell adhesion | GO:0007162 | 17% | 4.9E-10 | 3% | 0.08 |
| Granulocyte migration | GO:0097530 | 40% | 1.0E-24 | 19% | 1.7E-16 | Cell death | GO:0008219 | 9% | 1.5E-09 | 2% | 0.06 |
| Response to interferon-gamma | GO:0034341 | 40% | 1.0E-24 | 8% | 5.1E-05 | Cellular response to oxygen-containing compound | GO:1901701 | 10% | 3.0E-09 | 2% | 0.05 |
| Defense response to bacterium | GO:0042742 | 39% | 1.0E-24 | 14% | 1.7E-12 | Interleukin-12 production | GO:0032615 | 37% | 3.0E-09 | 2% | 0.57 |
| Myeloid leukocyte activation | GO:0002274 | 36% | 1.0E-24 | 12% | 6.8E-12 | Regulation of peptidyl-tyrosine phosphorylation | GO:0050730 | 18% | 3.6E-09 | 3% | 0.21 |
| Myeloid leukocyte migration | GO:0097529 | 35% | 1.0E-24 | 14% | 2.2E-16 | T cell mediated immunity | GO:0002456 | 26% | 4.7E-09 | 3% | 0.21 |
| Regulation of innate immune response | GO:0045088 | 34% | 1.0E-24 | 12% | 3.3E-12 | Positive regulation of lymphocyte differentiation | GO:0045621 | 27% | 5.5E-09 | 5% | 0.06 |
| Leukocyte chemotaxis | GO:0030595 | 33% | 1.0E-24 | 15% | 3.01E-17 | Regulation of interleukin-12 production | GO:0032655 | 37% | 1.4E-08 | 2% | 0.56 |

**Table 2.** Top 20 Gene Ontology molecular functions based on adjusted p-value for both sexes and females only

Cellular Components
| Both Sexes |  |  |  |  |  | Female Specific |  |  |  |  |  |
| --- | --- | --- | --- | --- | --- | --- | --- | --- | --- | --- | --- |
| Name | GO Identifier | Female %DEG of Total | Female adj-p | Male %DEG of Total | Male adj-p | Name | GO Identifier | Female %DEG of Total | Female adj-p | Male %DEG of Total | Male adj-p |
| Cell periphery | GO:0071944 | 9% | 1.0E-24 | 2% | 8.4E-10 | MHC protein complex | GO:0042611 | 61% | 5.8E-09 | 0% | - |
| Cell surface | GO:0009986 | 20% | 1.0E-24 | 5% | 2.0E-08 | **MHC class II protein complex** | GO:0042613 | 88% | 2.5E-07 | 0% | - |
| Plasma membrane | GO:0005886 | 10% | 1.0E-24 | 3% | 3.4E-11 | Plasma membrane protein complex | GO:0098797 | 13% | 3.1E-07 | 3% | 0.17 |
| Side of membrane | GO:0098552 | 22% | 1.0E-24 | 6% | 6.1E-10 | Receptor complex | GO:0043235 | 14% | 2.3E-06 | 2% | 0.35 |
| External side of plasma membrane | GO:0009897 | 29% | 1.0E-24 | 8% | 6.1E-10 | Inflammasome complex | GO:0061702 | 70% | 2.9E-06 | 10% | 0.43 |
| Membrane | GO:0016020 | 7% | 8.0E-22 | 2% | 9.0E-10 | Lytic vacuole | GO:0000323 | 11% | 6.8E-06 | 1% | 0.96 |
| Extracellular region | GO:0005576 | 11% | 3.5E-16 | 3% | 5.2E-03 | Lysosome | GO:0005764 | 11% | 6.8E-06 | 1% | 0.96 |
| Intrinsic component of membrane | GO:0031224 | 8% | 8.7E-16 | 2% | 4.8E-06 | Vesicle | GO:0031982 | 8% | 1.6E-05 | 2% | 0.57 |
| Extracellular space | GO:0005615 | 12% | 3.7E-14 | 3% | 7.8E-03 | Plasma membrane signaling receptor complex | GO:0098802 | 18% | 2.6E-05 | 4% | 0.25 |
| Integral component of membrane | GO:0016021 | 8% | 3.7E-14 | 2% | 4.9E-05 | Immunological synapse | GO:0001772 | 33% | 3.1E-05 | 10% | 0.06 |
| Phagocytic vesicle | GO:0045335 | 26% | 1.7E-10 | 6% | 2.4E-02 | Intracellular vesicle | GO:0097708 | 8% | 6.2E-05 | 2% | 0.56 |
| Integral component of plasma membrane | GO:0005887 | 11% | 3.4E-09 | 3% | 2.2E-03 | Cytoplasmic vesicle | GO:0031410 | 8% | 8.6E-05 | 2% | 0.55 |
| Intrinsic component of plasma membrane | GO:0031226 | 11% | 9.1E-09 | 3% | 3.7E-03 | NADPH oxidase complex | GO:0043020 | 71% | 1.3E-04 | 14% | 0.35 |
| Endocytic vesicle | GO:0030139 | 18% | 5.4E-08 | 5% | 1.9E-02 | Vacuole | GO:0005773 | 10% | 2.1E-04 | 1% | 1.00 |
| Membrane raft | GO:0045121 | 13% | 4.1E-06 | 4% | 6.3E-03 | IPAF inflammasome complex | GO:0072557 | 80% | 5.8E-04 | 0% | - |
| Membrane microdomain | GO:0098857 | 13% | 4.2E-06 | 4% | 6.3E-03 | AIM2 inflammasome complex | GO:0097169 | 80% | 5.8E-04 | 20% | 0.30 |
| Phagocytic cup | GO:0001891 | 40% | 6.8E-05 | 15% | 2.5E-02 | Phagocytic vesicle membrane | GO:0030670 | 35% | 6.7E-04 | 5% | 0.65 |
| Fc receptor complex | GO:0032997 | 100% | 2.3E-02 | 100% | 3.7E-03 | Lysosomal membrane | GO:0005765 | 14% | 1.0E-03 | 1% | 1.00 |
| Fc-epsilon receptor I complex | GO:0032998 | 100% | 2.3E-02 | 100% | 3.7E-03 | Lytic vacuole membrane | GO:0098852 | 14% | 1.0E-03 | 1% | 1.00 |
| Integrin alphaM-beta2 complex | GO:0034688 | 100% | 2.3E-02 | 100% | 3.7E-03 | Cellular anatomical entity | GO:0110165 | 5% | 1.1E-03 | 1% | 0.05 |

**Table 3.** Top 20 Gene Ontology cellular components based on adjusted p-value for both sexes and females only

Molecular Functions
| Both Sexes |  |  |  |  |  | Female Specific |  |  |  |  |  |
| --- | --- | --- | --- | --- | --- | --- | --- | --- | --- | --- | --- |
| Name | GO Identifier | Female %DEG of Total | Female adj-p | Male %DEG of Total | Male adj-p | Name | GO Identifier | Female %DEG of Total | Female adj-p | Male %DEG of Total | Male adj-p |
| Transmembrane signaling receptor activity | GO:0004888 | 19% | 4.9E-22 | 6% | 5.0E-08 | Cytokine binding | GO:0019955 | 24% | 2.9E-08 | 3% | 0.25 |
| Signaling receptor activity | GO:0038023 | 17% | 4.9E-22 | 5% | 1.3E-07 | *MHC protein complex binding* | GO:0023023 | 75% | 4.6E-08 | 17% | 0.06 |
| Molecular transducer activity | GO:0060089 | 17% | 4.9E-22 | 5% | 1.3E-07 | Antigen binding | GO:0003823 | 38% | 1.3E-07 | 3% | 0.52 |
| Immune receptor activity | GO:0140375 | 46% | 8.0E-22 | 18% | 9.8E-10 | *MHC class II protein complex binding* | GO:0023026 | 88% | 5.7E-07 | 0% | - |
| Signaling receptor binding | GO:0005102 | 12% | 9.9E-15 | 3% | 6.9E-05 | Protein binding | GO:0005515 | 6% | 7.7E-07 | 1% | 0.17 |
| Cytokine receptor activity | GO:0004896 | 36% | 1.9E-10 | 9% | 5.9E-03 | Peptide antigen binding | GO:0042605 | 50% | 4.2E-06 | 6% | 0.35 |
| Chemokine activity | GO:0008009 | 57% | 1.7E-09 | 30% | 4.6E-07 | Toll-like receptor binding | GO:0035325 | 57% | 5.4E-06 | 14% | 0.06 |
| Chemokine receptor binding | GO:0042379 | 42% | 3.0E-08 | 21% | 5.7E-06 | Pattern recognition receptor activity | GO:0038187 | 42% | 8.4E-05 | 5% | 0.36 |
| Signaling receptor regulator activity | GO:0030545 | 16% | 4.2E-08 | 5% | 1.4E-03 | CXCR3 chemokine receptor binding | GO:0048248 | 100% | 2.5E-04 | 25% | 0.15 |
| Cytokine activity | GO:0005125 | 25% | 7.8E-08 | 8% | 1.4E-03 | CARD domain binding | GO:0050700 | 46% | 7.4E-04 | 8% | 0.29 |
| Receptor ligand activity | GO:0048018 | 16% | 1.0E-06 | 5% | 6.4E-03 | Superoxide-generating NAD(P)H oxidase activity | GO:0016175 | 80% | 9.9E-04 | 0% | - |
| Immunoglobulin binding | GO:0019865 | 67% | 1.2E-06 | 50% | 2.0E-07 | Peptidase inhibitor activity | GO:0030414 | 18% | 9.9E-04 | 4% | 0.19 |
| Cytokine receptor binding | GO:0005126 | 17% | 1.6E-06 | 6% | 2.5E-03 | Peptidase regulator activity | GO:0061134 | 15% | 1.6E-03 | 4% | 0.15 |
| Signaling receptor activator activity | GO:0030546 | 15% | 1.8E-06 | 4% | 7.7E-03 | Endopeptidase inhibitor activity | GO:0004866 | 18% | 1.9E-03 | 4% | 0.17 |
| MHC protein binding | GO:0042287 | 38% | 4.7E-06 | 14% | 5.3E-03 | Lipopeptide binding | GO:0071723 | 67% | 2.4E-03 | 17% | 0.19 |
| 2'-5'-oligoadenylate synthetase activity | GO:0001730 | 86% | 5.9E-06 | 86% | 4.6E-09 | T cell receptor binding | GO:0042608 | 38% | 2.4E-03 | 0% | - |
| Immunoglobulin receptor activity | GO:0019763 | 100% | 1.7E-05 | 60% | 4.3E-04 | Acetylcholine receptor inhibitor activity | GO:0030550 | 67% | 2.4E-03 | 17% | 0.19 |
| Molecular function regulator | GO:0098772 | 8% | 2.7E-05 | 2% | 2.1E-02 | Lipid binding | GO:0008289 | 9% | 3.3E-03 | 2% | 0.25 |
| MHC class I protein binding | GO:0042288 | 38% | 6.6E-05 | 17% | 3.0E-03 | TAP2 binding | GO:0046979 | 42% | 4.3E-03 | 8% | 0.28 |
| IgG binding | GO:0019864 | 83% | 8.4E-05 | 50% | 7.71E-04 | CXCR chemokine receptor binding | GO:0045236 | 57% | 4.3E-03 | 14% | 0.21 |

### Blockade of MHC II attenuates muscle hyperalgesia in female but not male mice

To further explore the sex differences in MHC II signaling, we blocked MHC II during induction of muscle pain to determine if it would prevent muscle hyperalgesia in female mice (Fig. 3A). On both the ipsilateral and contralateral limb, blockade of MHC II resulted in a significant effect for group (Ipsilateral: F_1,27_=15.19, p<0.01; Contralateral: F_1,27_=20.46, p<0.01) and group by sex (Ipsilateral: F_1,27_=4.38, p=0.04; Contralateral: F_1,27_=13.07, p<0.01) for 24 hour MWT values which justified the separation of data by sex. Blockade of MHC II attenuated the development of muscle hyperalgesia in females seen on the ipsilateral (MHC II Ab: 1250.17±64.1; Control: 998.0±17.89; t_14_=3.78, p<0.01, Cohen’s d=1.89) and contralateral limb (MHC II Ab: 1271.08±52.52; Control:1018.12±10.67; t14=4.72, p<0.01; Cohen’s d=2.36) 24 hours after induction of the model (Fig. 3B). However, blockade of MHC II had no impact on the development of muscle hyperalgesia in male mice on the ipsilateral limb 24 hours after induction of the model (MHC II Ab: 1114.29±46.59; Control:1041.0±22.36; t_14_=1.42, p=0.18; Cohen’s d=0.71) (Fig. 3C). We also calculated MWT change scores between baseline and 24 hour MWT values. In females, there was a significant difference for MWT change score values on the ipsilateral (MHC II Ab: 361.45±74.75; Control: 617.33±26.66; t_14_=3.22, p<0.01) and contralateral (MHC II Ab: 350.87±52.04; Control: 617.21±23.29; t14=4.67, p<0.01) limb (Fig. 3D). Again, blockade of MHC II resulted in no effect in males on the ipsilateral limb (MHC II Ab: 460.33±42.53; Control: 546.58±38.26; t_14_=1.51, p=0.15) (Fig. 3E). This further confirms the female specific activation of MHC II signaling and its role in the production of muscle hyperalgesia following induction of activity-induced muscle pain.

**Figure 3.**
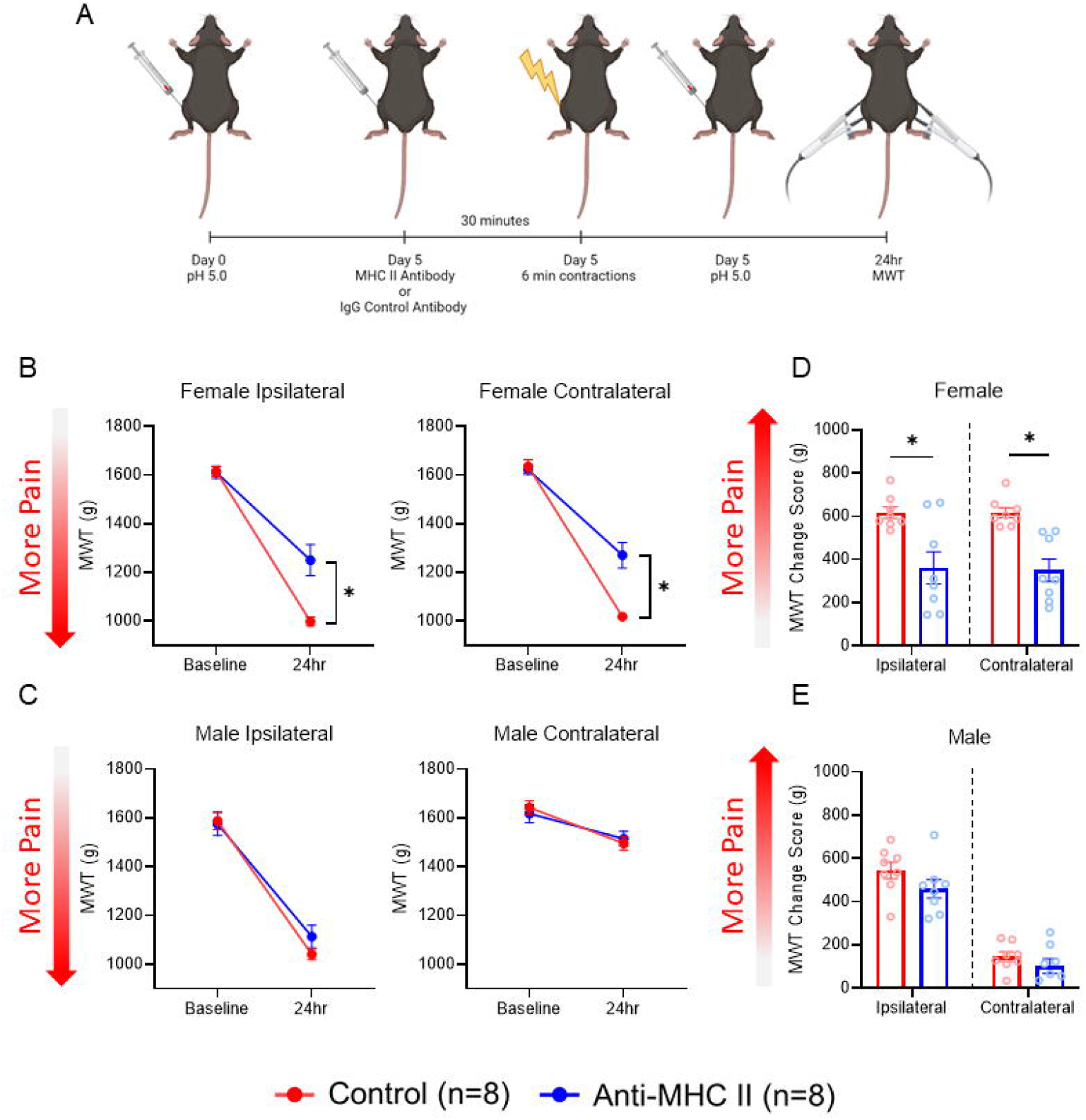
Blockade of MHC II attenuates development of muscle hyperalgesia in female but not male mice. A. Graphical depiction of experimental protocol to block MHC II during induction of muscle pain. B. Muscle withdrawal thresholds (MWT) for female mice. Blockade of MHC II resulted in attenuation in the reduction of MWT at the 24 hour time point when compared with animals who received the control antibody on both the ipsilateral and contralateral limb. C. MWT for male mice. Blockade of MHC II did not alter the development of muscle hyperalgesia on the ipsilateral limb. D. Change scores for MWT values for female mice. There was a greater reduction in MWT values in females who received the control antibody when compared with animals who received MHC II antibody on both the ipsilateral and contralateral limb. E. Change scores for MWT values for male mice. There was no difference in MWT change scores between males who received the control or MHC II antibody on the ipsilateral limb. *p<0.05 compared with control antibody; Data are mean±SEM; Images made on BioRender.

### Physical activity prevents activity-induced muscle pain through blunting of immune response

Prior studies show 8 weeks of prior physical activity prevents the onset of muscle pain (*17, 18, 20, 28*). To further explore potential mechanisms for how physical activity prevents development of activity-induced muscle hyperalgesia, we utilized RNA sequencing of the left gastrocnemius muscle from physically active mice and compared results to sedentary mice and pain free controls. Surprisingly, in males, between sedentary and physically active animals who received the pain model there were only 11 DEGs (9 upregulated, 2 downregulated) of which most upregulated genes were myosin or troponin genes which encode for skeletal muscle protein. In males, between physically active mice who received the pain model and pain free controls, there were 111 DEGs (105 upregulated, 6 downregulated). In females, between sedentary and physically active animals who received the pain model, there were 334 DEGs (331 upregulated, 3 downregulated) (Fig. 4A Left Panel) and 69 DEGs (60 upregulated, 9 downregulated) between physically active mice who received the pain model and pain free controls (Fig. 4A Right Panel).

**Figure 4.**
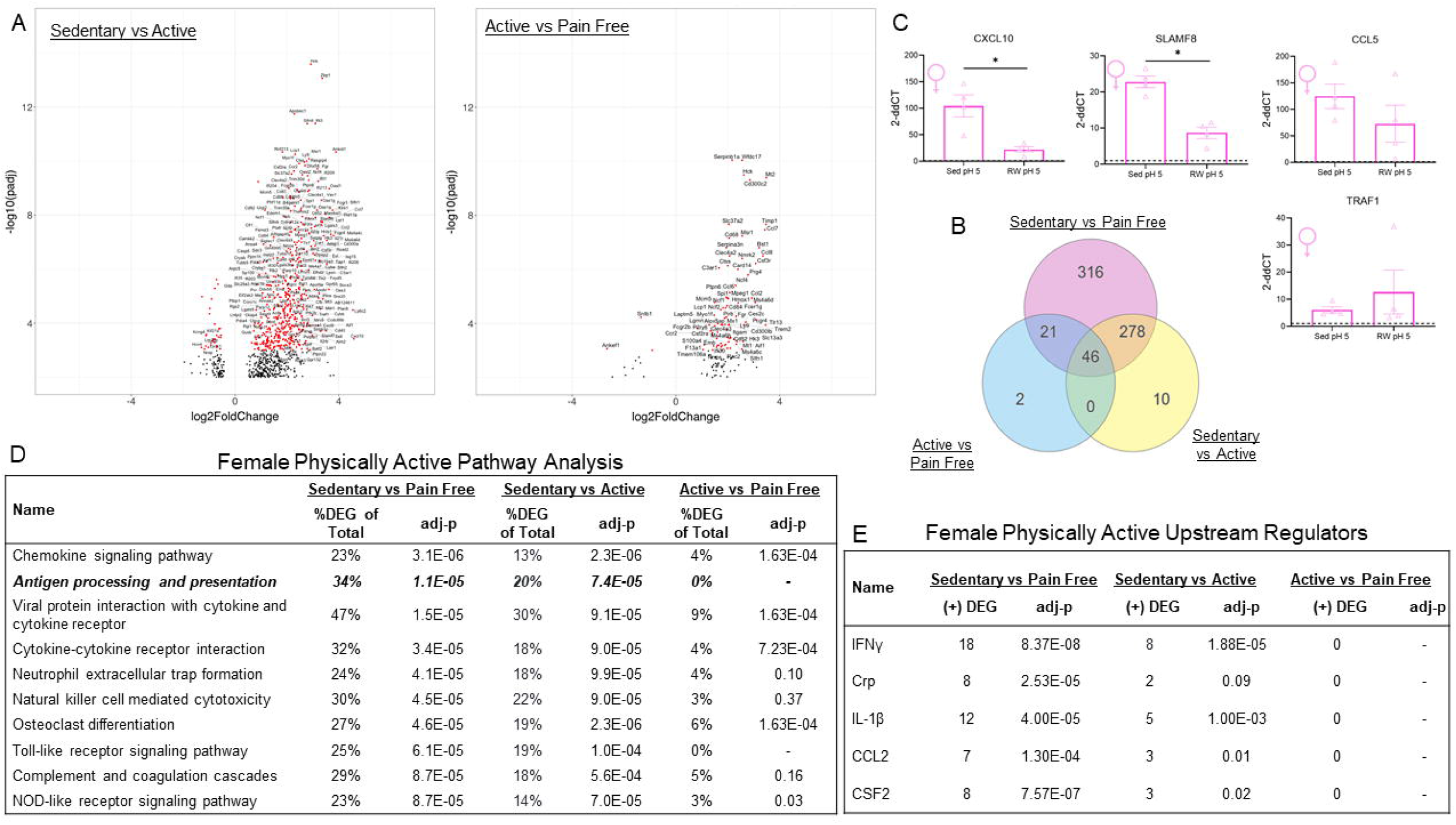
Prior physical activity blunts enrichment of several immune system pathways including a complete prevention of activation of antigen processing and presentation in females. A. Volcano plots of differentially expressed genes (DEGs) in females between sedentary and physically active animals who received the muscle pain model (left) and between physically active animals who received the muscle pain model and pain free controls (right). Only DEG’s with an adjusted p-value <0.1 are plotted with red dots depicting a more significant p-value (adj-p:<0.001). B. Venn Diagram depicting DEG between the 3 treatment conditions. Of note, 278 genes were both differentially expressed between sedentary and pain free animals and between sedentary and physically active animals but not between physically active and pain free controls. This suggests physical activity worked to prevent the transcriptional changes of these 278 genes in response to induction of muscle pain. C. qPCR confirmation of select genes from RNA sequencing analysis. Physical activity prevented increases in *CXCL10* and *SLAMF8* but not *CCL5* and *TRAF1*. Dotted line represents expression level for pain free controls. D. Top 10 enriched pathways with smallest adjusted p-values from RNA sequencing analysis for female mice between sedentary and physically active who received the activity-induced muscle pain model and pain free controls. Prior physical activity prevented enrichment of numerous pathways seen as a decrease in the % of DEGs upregulated in that pathway. Prior physically activity completely blocked the activation of several immune system pathways including antigen processing and presentation. E. Upstream regulators predicted to be present based on DEGs between the three treatment groups. The predicted presence of these upstream regulators, including IFNγ, was prevented by prior physical activity. *p<0.05 compared with sedentary pH 5.0; Sed=Sedentary, RW=Running Wheel, IL=interleukin, IFNγ=interferon gamma, Crp= C reactive protein, CSF= colony stimulating factor; Data are mean+SEM.

To explore potential mechanisms by which physical activity prevents muscle pain we performed a pathway analysis of all 3 treatment groups. Since males had few DEGs between sedentary and physically active mice, we were unable to perform a 3-way analysis and thus we only analyzed female data. A 3-way meta-analysis of female DEGs revealed a set of 278 DEGs that were upregulated between sedentary and pain free controls and also between sedentary and physically active mice but not between physically active and pain free controls. This suggests physical activity prior to induction of the pain model blocked the upregulation of these 278 genes (Fig 4B).

To confirm RNA sequencing results, 4 genes were selected, and expression levels were measured with qPCR in a separate set of animals within the same experimental groups. Animals with prior physical activity had lower expression levels of *CXCL10* (t_6_=3.82, p<0.01) and *SLAMF8* (t_6_=6.23, p<0.01) compared with sedentary animals (Fig 4C). There were no significant differences for *CCL5* (t_6_=1.24, p=0.26) or *TRAF1* (t_6_=0.80, p=0.45) between physically active and sedentary animals (Fig 4C). Thus, we were able to validate 2 out of 4 genes from our RNA sequencing analysis in which physical activity prevented their upregulation.

Pathway analysis and upstream regulators revealed that prior physical activity decreased the upregulation of genes in several immune system pathways demonstrated by the difference in number of DEGs in the sedentary versus pain free compared with the physically active versus pain free animals (Fig. 4D). Of note, prior physical activity prevented activation of several pathways including antigen processing and presentation, neutrophil extracellular trap formation, natural killer cell mediated cytotoxicity, toll-like receptor signaling pathway, and compliment and coagulation cascades. Upstream regulators also showed that physical activity prevented upregulation of genes associated with the presence of several cytokines including INFγ and IL-1β (Fig. 4E) This suggests that prior physical activity dampens activation of immune system pathways to protect against development of muscle pain in females.

### Activity-induced muscle pain increases macrophages in muscle and their phenotype is altered by prior physical activity

To further explore alterations in the immune system and MHC II signaling in response to the induction of activity-induced muscle pain, we performed flow cytometry on the left gastrocnemius muscle 24 hours after induction of the pain model in sedentary, physically active and pain free controls. Unless otherwise noted, no significant group by sex interaction was determined for flow cytometric data and therefore male and female data were analyzed together. Gating strategy for flow cytometry experiments are located in Figure 5A. After induction of the pain model, there was a greater percentage of immune cells (% of CD45+) in the muscle in both sedentary and physically active mice when compared with pain free controls (F_2,18_=30.21, p<0.01; 7.2 vs Sed 5 p<0.01, 7.2 vs Ex 5 p<0.01) (Fig. 5B). This increase in total immune cells was driven predominately by a higher percentage of immune cells that were Ly6C+ (CD45+CD3-LyC6+) as both sedentary and physically active animals had higher percentages of Ly6C+ cells compared with pain free controls (F_2,18_=19.78, p<0.01; 7.2 vs Sed 5 p<0.01, 7.2 vs Ex 5 p<0.01) (Fig. 5C+D). The majority of Ly6C+ cells (75-84%) were macrophages (F4/80+) in all three groups. There was a significant effect for treatment group for total number of macrophages (F_2,18_=82.69, p<0.01) with both sedentary and physically active mice that received the pain model showing a greater number of macrophages when compared with pain free animals (p<0.01), and physically active mice showing a greater number of macrophages when compared with sedentary animals (p<0.01) (Fig. 5E+F). Sedentary animals also had a higher number of CD3+ cells when compared with physically active and pain free controls (F_2_,_18_=57.99, p<0.01; 7.2 vs Sed 5 p<0.01, Sed 5 vs Ex 5 p<0.01) (Fig. 5G). Due to the low number of cells (<1000 events) we were not able to analyze group differences for T cell subsets (CD4, CD8, CD183, CD184, CD25, CD103, CD196, CD69), antigen presenting cell costimulatory molecule CD80, natural killer cells (CD49b) or dendritic cells (CD205).

**Figure 5.**
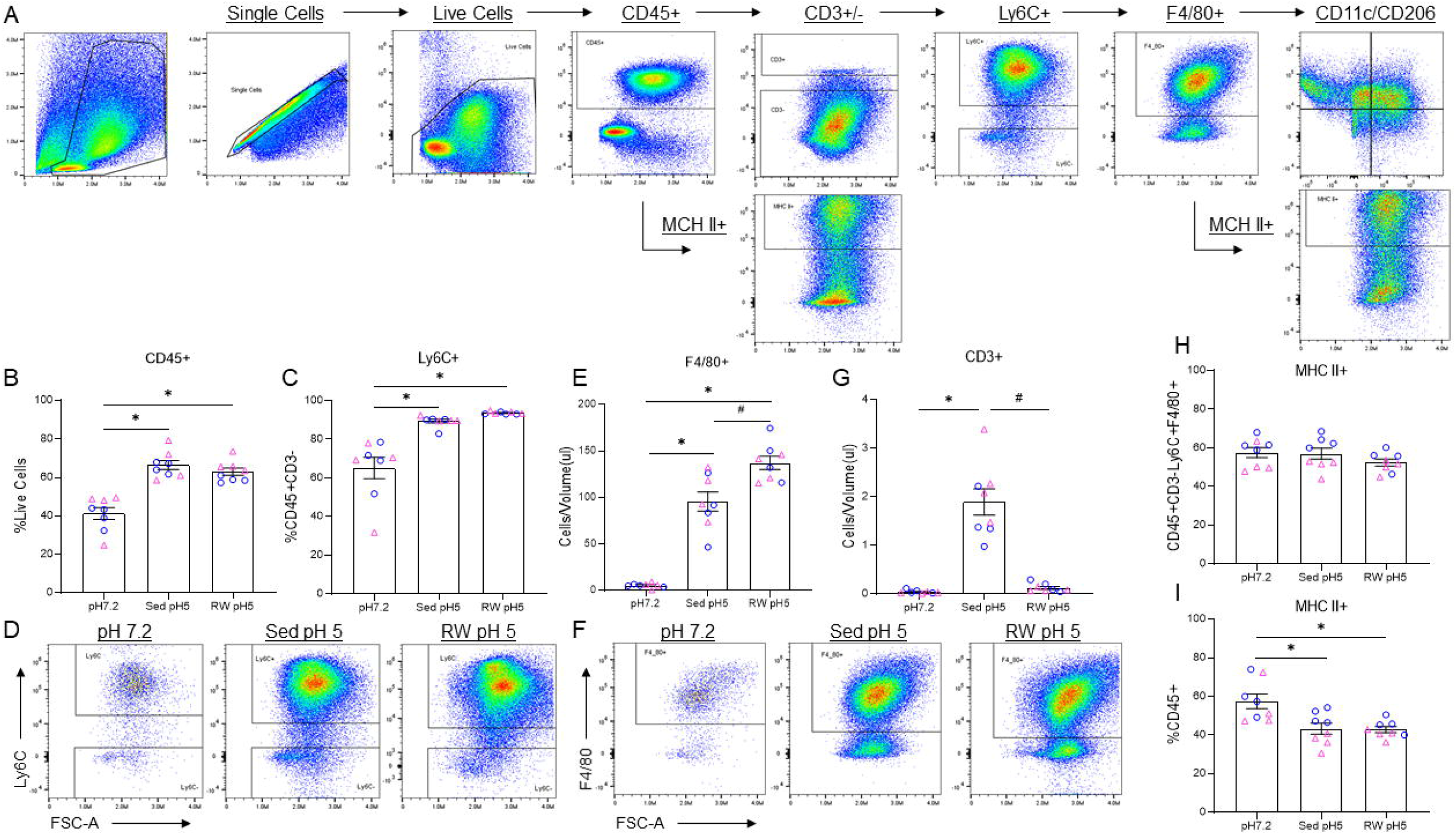
Activity-induced muscle pain alters immune cell populations in the gastrocnemius muscle in both males and females. A. Depiction of gating strategy for flow cytometry analysis. B. Sedentary and physically active animals who received muscle pain model had significantly higher percentage of CD45+ cells in the gastrocnemius muscle compared with pain free controls. C. Sedentary and physically active animals who received muscle pain had a significantly higher percentage of CD45+ cells that were Ly6C+ when compared with pain free controls. D. Representative flow cytometry data for Ly6C+ cells for pain free and sedentary and physically active animals who received the pain model. E. The muscles of sedentary and physically active animals who received muscle pain had a significantly higher number of macrophages (CD45+CD3-Ly6C+F4/80+) when compared with pain free controls. Physically active animals also had a significantly higher number of F4/80+ cells compared with sedentary mice. F. Representative flow cytometry data for F4/80+ cells for pain free and sedentary and physically active animals who received the pain model. G. Sedentary animals had a significantly higher amount of CD3+ cells following induction of muscle pain when compared with physically active animals and pain free controls. H. There were no differences in percentage of macrophages expressing MHC II between the treatment groups. I. There was a significant group difference in the percentage of immune cells expressing MHC II with sedentary and physically active animals having fewer than pain free controls. *p<0.05 compared with sedentary, pH 7.2, #p<0.05 compared with Sed pH 5; Sed=Sedentary, RW=Running Wheel; Blue circles represent male data, Pink triangles represent female data; Data are mean+SEM.

MHC II is found on antigen presenting cells including macrophages (*45*). Since we saw an increase in MHC II signaling with RNA sequencing analysis, and macrophages are the predominate cell type in the muscle, we tested if macrophages had increased MHC II expression in the muscle following induction of muscle pain. However, there were no differences between groups in the number of MHC II+ positive macrophages (F4/80+MHC II+) (F_2,18_=2.7, p=0.09) (Fig. 5H). It is possible that MHC II could be increased on other immune cells (dendritic cells, B cells) (*45*), therefore, we explored if there were differences in the number of total immune cells expressing MHC II+ (CD45+MHCII+). There were significantly fewer number of CD45+ immune cells expressing MHC II with sedentary and physically active mice expressing less MHC II+ compared with control mice (F_2,18_=10.66, p<0.01; 7.2 vs Sed 5 p<0.01, 7.2 vs Ex 5 p<0.01) (Fig. 5I). This lower amount of MHC II expressing immune cells is likely due to the increased total of CD45+ cells seen in the animals who received the muscle pain model.

To further explore alterations found in macrophages, we analyzed differences in macrophage phenotype in the gastrocnemius muscle by examining cells expressing an M1 or M2 marker (Fig. 6A). There were significant differences in macrophage phenotype for M1 expression (CD206-,CD11c+) (F_2,18_=627.86, p<0.01), M1/2 co-expression (CD206+,CD11c+) (F_2,18_=178.16, p<0.01), M2 expression (CD206+,CD11c-) (F_2,18_=465.87, p<0.01) and undifferentiated macrophages M0 (CD206-,CD11c-) (F_2,18_=252.45, p<0.01). Male and female data were analyzed separately for the M0 phenotype due to a significant sex by group interaction (F_2,18_=4.09, p=0.03). Sedentary mice with the pain model had a higher percentage of macrophages that were M1 or M1/M2 phenotype compared with pain free and physically active animals (p<0.01) (Fig 6B+C). Physically active animals had a higher percentage of M2 macrophages compared with sedentary animals (p<0.01), and M0 phenotype compared with sedentary and pain free animals (Male p<0.01; Female p<0.01) (Fig. 6D+E). Pain free animals had a higher percentage of macrophages that were M2 phenotype compared with sedentary and physically active animals (p<0.01) (Fig. 6E). This suggests prior physical activity could be preventing the development of muscle pain by stopping the polarization of macrophages to an M1 phenotype after muscle insult.

**Figure 6.**
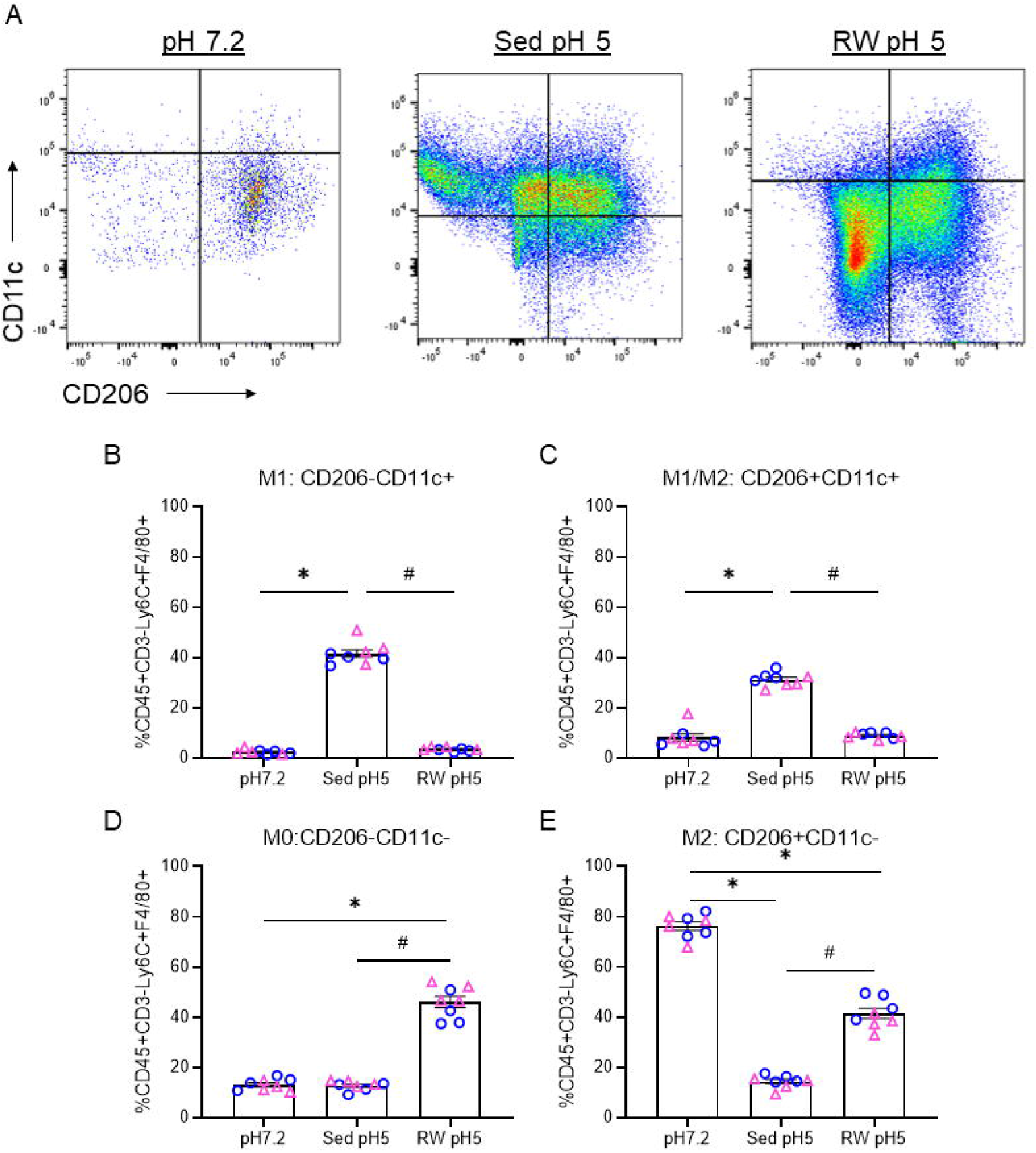
Prior physical activity impacts the phenotype of macrophages in gastrocnemius muscle after induction of activity-induced muscle pain in both males and females. A. Representative image of flow cytometry data for macrophage phenotype in pain free and sedentary and physically active animals who received the pain model. B. Sedentary animals had a higher percentage of macrophages that displayed the M1 phenotype when compared with both physically active and pain free controls after induction of muscle pain. C. Sedentary animals also had a higher percentage of macrophages that displayed the M1/2 phenotype when compared with both physically active and pain free controls after induction of muscle pain. D. Physically active animals had a higher percentage of macrophages that displayed the M0 phenotype when compared with sedentary and pain free animals. E. Pain free animals had a higher percentage of macrophages that displayed an M2 phenotype compared with both sedentary and physically active animals who received the muscle pain model. Physically active animals had a higher percentage of macrophages that displayed the M2 phenotype when compared with sedentary animals. *p<0.05 compared with sedentary pH 7.2, #p<0.05 compared with RW pH 5; Sed=Sedentary, RW=Running Wheel; Blue circles represent male data, Pink triangles represent female data; Data are mean+SEM.

### Modulation of Testosterone

We previously reported that the sex hormone testosterone mediates the sex-dependent pain phenotype produced by the activity-induced muscle pain model (*25*). To determine if testosterone impacts activation of DEGs, we manipulated circulating levels of testosterone in male and female mice prior to induction of activity-induced muscle pain and examined mRNA expression in female-specific genes (*KLRK1, CD74, H2-Aa, and MILR1*) (Fig. 7A). In males, orchiectomy to remove testosterone had no effect on expression of *KLRK1, CD74, H2-Aa*, and *MILR1* (t_10_=1.26-1.93, p=0.08-0.23) when compared to sham surgery after induction of muscle pain (Fig. 7B). This suggests that modulating testosterone in males does not alter expression of some female specific immune system genes.

**Figure 7.**
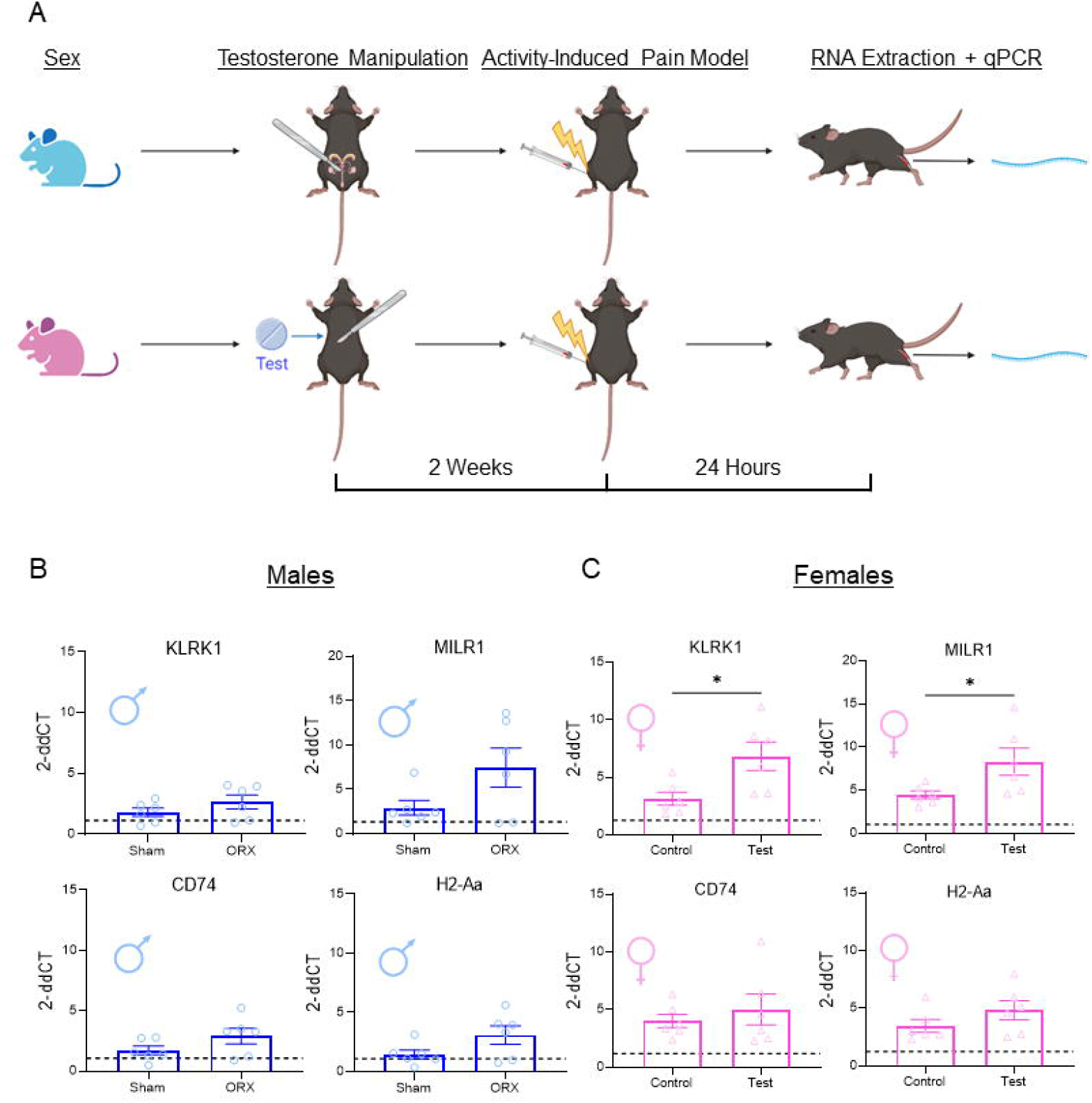
Modulation of testosterone prior to induction of muscle pain does not alter female DEG’s in males and only some genes in females. A. Graphical depiction of experimental design to modulate testosterone prior to induction of activity-induced muscle pain, for male and female mice. B. In males, decreasing testosterone through orchiectomy prior to induction of muscle pain did alter the expression levels of female specific DEG’s when compared with sham operated males after induction of muscle pain. C. In females, increasing testosterone through testosterone pellet implantation prior to induction of the muscle pain model caused a higher expression of *KLRK1* and *MILR1* when compared with animals who received control pellets after induction of muscle pain. Dotted line represents expression level for pain free controls. *p<0.05 compared with controls; ORX=orchiectomy, Test=tesosterone; Data are mean+SEM. Images made on BioRender.

In females, increasing testosterone systemically, resulted in higher expression of *KLRK1* and *MILR1* (t_10_=2.35-2.72, p=0.02-0.04), but not *CD74 and H2-A* (t_10_=0.68-1.35, p=0.20-0.51), when compared to animals who received control pellets after induction of muscle pain (Fig. 7C). This suggests that testosterone may have a partial impact on immune system activation in females.

## Discussion

### Overview

Using an unbiased RNA sequencing approach, the current study showed activation of the immune system with a robust sex-specific effect in females. Specifically, females showed activation of the antigen processing and presentation pathway along with MHC II signaling, which was confirmed in separate animals with qPCR. Further, blockade of MHC II during induction of muscle pain, attenuated development of muscle hyperalgesia in female, but not male mice, confirming a sex-specific role for this pathway in development of muscle pain. Regular physical activity attenuated the immune response in female mice as measured with RNA sequencing. We further showed an increase in the number of macrophages and T cells in the muscle after induction of muscle pain, with sedentary animals expressing a larger percentage of M1 macrophages while physically active and pain free animals expressing a larger proportion of M2 and M0 macrophages. Thus, we uncovered alterations in local immune mechanisms after induction of muscle pain in both male and female mice, with additional sex-specific mechanisms activated only in females.

### Sex differences and MHC II signaling

The main goal of this project was to explore potential sex-specific transcriptional alterations induced by muscle pain in sedentary and physically active mice. Sedentary females, compared to sedentary males, showed a greater number and magnitude of change in DEGs after muscle insult. While several pathways were increased in both sexes, more DEGs were upregulated in sedentary females across multiple immune pathways including chemokine signaling, viral protein interaction, toll-like receptor signaling, complement and coagulation cascades, and NOD-like receptor signaling which prior literature has shown are important for the generation of pain in several animal models (*46–50*). Further, a role for IL-1β and IL-33 were implicated in both sexes which is consistent with prior research in animal models of pain (*14, 51*). In fact, our prior research using this animal model shows blockade of IL-1β prevents development of chronic pain in male and female mice (*14*). This data suggests that production of muscle pain in male and female mice involves activation of several immune system cells and pathways and is not restricted to a single cell type or mechanism.

The most robust sex difference was a female-specific activation of the antigen processing and presentation pathway particularly involving MHC II signaling. The antigen processing and presentation pathway is a function of the adaptive immune system and is mediated through MHC I and MHC II molecules on antigen presenting cells (*45*). MHC I molecules are present on nearly all nucleated cells and communicate with CD8+ T cells in response to an endogenous antigen (i.e. viral infection) (*45*). MHC II molecules are present on antigen presenting cells (macrophages, B cells, dendritic cells) and communicate with CD4+ T cells in response to phagocytosis of an exogenous antigen (*45*). Interestingly, the current study discovered a female exclusive increase in MHC II signaling using RNA sequencing in muscle tissue following induction of muscle pain, and pharmacological blockade of MHC II prevented muscle hyperalgesia in female, but not male mice. Consistent with our findings, MHC II expression is increased on macrophages in the hind paw following induction of a prolactin pain model in female, but not male, mice (*8*). Similarly, in studies utilizing only female mice MHC II protein is increased in the spinal cord in bone cancer pain (*52*) and on Schwann cells at site of injury in neuropathic pain (*53, 54*), and genetic deletion of MHC II in Schwann cells prevents hyperalgesia in neuropathic pain (*53*). In contrast, prior studies using only male animals show MHC II protein is increased in the spinal cord and dorsal root ganglia (DRG) in neuropathic pain models (*55–60*), and pharmacological blockade of MHC II (*57, 58*) or genetic deletion of MHC II reduces pain behaviors in an animal model of nerve injury (*56*). Thus, a lack of prior studies utilizing both sexes make it difficult to determine if there is a sex difference in MHC II signaling in the context of pain. It is likely that the female specific increases in MHC II signaling is species, location, and pain model specific. While we showed an increase in MHC II signaling with RNA sequencing, we did not find an increase of MHC II protein on immune cells with flow cytometry. It is likely that protein increases occur after increases in mRNA. Indeed, prior studies show increases in MHC II protein in DRG and spinal cord of animals 7-10 days following induction of neuropathic pain (*55, 56, 61*). Future work will need to study protein analysis of MHC II in the muscle at later time points following induction of muscle pain.

Clinical studies also show increases in MHC II signaling in individuals with chronic pain. In humans, MHC genes are referred to as human leukocyte antigen (HLA) genes with HLA Class I (HLA-A, -B, -C) being synonymous to MHC I and HLA Class II (HLA-DR, -DP, -DQ) being synonymous with MHC II (*62, 63*). Increases in HLA Class II signaling are found in several chronic pain conditions including complex regional pain syndrome (*64–66*), rheumatoid arthritis (*67*), inflammatory bowel disease (*68, 69*), neuropathic pain (*70*), low back pain (*71*), and temporomandibular joint disorder (*72*). However, most of this research did not explore or was not adequately powered to study sex differences in HLA Class II signaling. One paper that did explore sex differences using RNA sequencing found a female specific increase in HLA Class II signaling in DRGs from individuals with neuropathic pain (*70*); however, it is unclear what types of cells in the DRGs were expressing HLA class II. Future work across various pain conditions needs to examine both sexes to adequately determine if increases are female specific and determine the cell types expressing HLA Class II.

### Muscle pain and immune cells

The current study shows the majority of immune cells in skeletal muscle are macrophages in both uninjured mice and after induction of muscle pain which agrees with prior studies using animal models of muscular dystrophy (*70, 73*). Similar to our prior work (*11*), the current study also showed an increase in the number of macrophages 24 hours after induction of the model. Macrophages play a critical role in development of muscle hyperalgesia as depletion of muscle macrophages and elimination of P2X4 receptors on muscle macrophages prevents hyperalgesia (*11–14*). Macrophages are plastic and change their phenotype in response to the external environment; the current study shows an increase in proportion of M1 macrophages in the muscle after induction of muscle pain in sedentary mice. This is in agreement with prior work showing an increase in M1 macrophages in the muscles of sedentary mice following injection of carrageenan (*13*). M1 macrophages release pro-inflammatory cytokines while M2 release antiinflammatory cytokines. Pro-inflammatory cytokines are important for induction of hyperalgesia. For example, injection of the inflammatory cytokine IL-6 in muscle produces hyperalgesia (*11*) while blockade of IL-1β prevents the development of muscle pain (*14*) in both male and female mice. On the other hand, blockade of the P2X7 pathway in muscle prevents development of muscle hyperalgesia only in male mice. Uniquely, the current study shows a sex-specific mechanism with female mice activating the antigen processing and presentation pathway with MHC II signaling to mediate muscle hyperalgesia. Thus, while there are common pain facilitating functions for specific immune cells and pathways between male and female mice, there are also sex-specific mechanisms that mediate the development of muscle hyperalgesia. The complexity of these mechanisms is likely critical to gaining a full understanding of chronic pain in both male and female mice and developing effective treatments that are sex specific.

Similar to prior work, we found little to no T-cells in the muscles of uninjured mice (*73, 74*). After muscle injury, however there was a small increase in the number of T-cells (CD3+) in both male and female sedentary mice. In mouse models of muscular dystrophy, increases in T-cells are present in the muscle 3-4 days after insult and last for 30 days (*75, 76*). Similarly, infiltrating T-cells peak at the site of insult 2-3 weeks after nerve injury (*77*). We propose that increases in MHC II signaling, from macrophages in muscle, could signal and activate CD4+ T-cells to contribute to the generation of muscle pain in females. A female specific role for T-cells in the spinal cord in the generation of neuropathic pain has been suggested (*9*). However, others show male mice and rats lacking T-cells are protected against development of neuropathic pain (*10, 56, 78*). These differences in findings could be due to the utilization of different species, strains, pain models, or timing after injury. It is also possible that tissue-specific differences, such as muscle, nerve, or spinal cord will show unique differences in the role of T-cells in the generation of pain.

### Impact of prior physical activity on immune mechanisms

The current study, using RNA sequencing, shows that prior physical activity attenuates the immune response to muscle insult, including the antigen processing and presentation pathway, in females. The current study also showed prior physical activity results in a higher proportion of macrophages demonstrating the M2 and M0 phenotype, and a lower proportion of the M1 phenotype, in the gastrocnemius muscle after induction of the pain model in both males and females. This agrees with prior work which showed increases in the M2 macrophage phenotype in muscle from uninjured animals and increases in M2 and decreases in M1 phenotypes in the muscle or at the site of nerve injury after exercise (*13, 15, 17, 79*). The higher proportion of antiinflammatory M2 macrophages could account for the attenuated immune response observed with RNA sequencing since M2 macrophages secrete anti-inflammatory cytokines such as IL-10 which protects against development of pain in several animal models (*13, 15, 17, 80–83*). For example, increasing expression of IL-10 in the knee of canines with osteoarthritis reduces veterinary and owner pain assessments (*80*). We previously demonstrate that blockade of anti-inflammatory cytokines IL-10 or IL-4 prevents the analgesia produced by regular physical activity (*17, 84*), and genetic deletion of IL-4 prevents analgesia and the increase in macrophage M2 phenotype produced by exercise (*15*). In addition, the current study shows the expansion of T-cell population in the muscle after tissue insult does not occur in physically active mice. IL-10 inhibits MHC II antigen presentation on immune cells to T-cells and thus could explain how physical activity prevents MHC II signaling and T-cell increases (*85*). Together these data suggest that regular physical activity dampens immune transcriptional changes and polarization of macrophages to a pro-inflammatory phenotype in response to a muscle insult.

Surprisingly, in males, prior physical activity did not prevent the increases in the immune response induced by muscle pain but did prevent increases in several muscle protein genes including troponin and myosin signaling. Increased expression of troponin and myosin muscle protein genes suggests the presence of muscle damage or fatigue due to the electrically stimulated muscle contractions in sedentary but not physically active mice (*86*). While we cannot explain the lack of effect on the immune changes, it is noted that physical activity prevents pain equally in both sexes (*17–19, 26, 28, 87*), and prior studies examining macrophage phenotype and anti-inflammatory cytokine contributions to exercise-induced analgesia were done primarily in males (*15, 17*). It is possible in males the analgesia produced by physical activity is a result of the altered phenotype of macrophages that results in enhanced secretion of anti-inflammatory cytokines, which is independent of changes in gene transcription. While females on the other hand showed a unique increase in the antigen processing and presentation pathway at the transcription level that was blocked by physical activity, and thus there are likely multiple mechanisms within and between sexes that mediate protection against muscle pain.

### Impact of testosterone on immune cell activation

We have previously shown that testosterone mediates the sex-dependent pain phenotype which our activity-induced muscle pain model produces (*25*). To test if testosterone levels have any impact on the female specific upregulated DEG’s in the muscle, we tested if manipulation of testosterone prior to induction of the pain model altered their expression levels in both sexes. In males, decreasing testosterone failed to increase the expression level of female specific DEG’s suggesting testosterone levels had no impact on the lack of upregulation of these genes. In females, increasing testosterone had the opposite effect of our hypothesized outcome as it increased the expression of *KLRK1 and MILR1* but had no effect on the expression levels of *CD74 and H2-Aa*. This suggests testosterone levels might have an impact on the activation of some but not all female specific DEG’s in female mice. Prior work shows a sex and testosterone dependent activation of specific immune cells in both the innate and adaptive immune system. Under normal conditions, adult males and females have different populations of circulating immune cells with females expressing a greater proportion of CD4+ T-cells and males expressing a greater proportion of natural killer cells and CD8+ T-cells (*88*). Following an immune insult, females demonstrate a larger immune response and have a greater proportion of both CD4+ and CD8+ T-cells (*88, 89*). The current study similarly shows a larger change in the immune system responses observed with RNA sequencing, and thus is consistent with these prior studies. Testosterone has been shown to modulate immune cell response to infectious insults. Orchiectomized male mice have increases in CD4+ and CD8+ T-cells and an enhanced immune response to infection when compared with intact males (*90, 91*). Female mice secrete more IFNγ compared with males in response to infection which is prevented by testosterone treatment (*92*). Despite these differences, testosterone had no effect on modulating specific DEGs in males and only partially modulated genes in females. Sex differences in immune system activation could be caused by other factors such as genetics and levels of estrogen and progesterone (*93, 94*).

## Conclusion

In summary, we used an unbiased approach to discover mechanisms that are enriched in skeletal muscle at the site of insult following induction of muscle pain in male and female mice. We discovered the majority of changes occurred in the immune system in both sexes, and there was a robust sex difference in which females showed activation of the antigen processing and presentation pathway with MHC II signaling. Our data show there are common changes across sexes in the transcriptome and immune cell populations, but there are also unique changes in females after muscle insult. We also show that prior physical activity alters how the local immune system responds to a muscle insult in terms of both transcriptional changes and macrophage phenotype polarization. This suggests that activation of immune pathways to a muscle insult are plastic and factors such as sex or level of physical activity can alter how the local immune system responds.

## Acknowledgments

The data presented herein were obtained at the Flow Cytometry Facility, which is a Carver College of Medicine / Holden Comprehensive Cancer Center core research facility at the University of Iowa. The facility is funded through user fees and the generous financial support of the Carver College of Medicine, Holden Comprehensive Cancer Center, and Iowa City Veteran’s Administration Medical Center.

## Supplemental Table Legends

Table 1 – Average distance run (km) per week for physically active animals for each experiment

Table 2 – Accession number and forward and reverse sequences used for qPCR primers

Table 3 – Antibodies and their working concentrations used for flow cytometry experiment

## Supplemental Figure Legends

Supplemental Figure 1

**Principal component analysis for top 500 variable genes from RNA sequencing analysis**

PCA results reveal that 68.92% of the variance is explained by PC1 and 9.47% of the variance is explained by PC2. A. Samples colored by group assignment demonstrate animals cluster together based on group. B. Samples colored by sex demonstrate animals cluster together based on sex. C. Samples colored by date of tissue pull demonstrate lack of clustering suggesting no batch effect in RNA sequencing results.

Supplemental Figure 2

**Sample-to-sample distance heatmap for RNA sequencing results**

Samples distance compared based on top 500 variable genes from RNA sequencing results.

The greater the distance is shown in dark red and the lesser the distance is shown in dark blue.

GM-45 displayed great distance from other samples and was removed from RNA sequencing analysis.

---

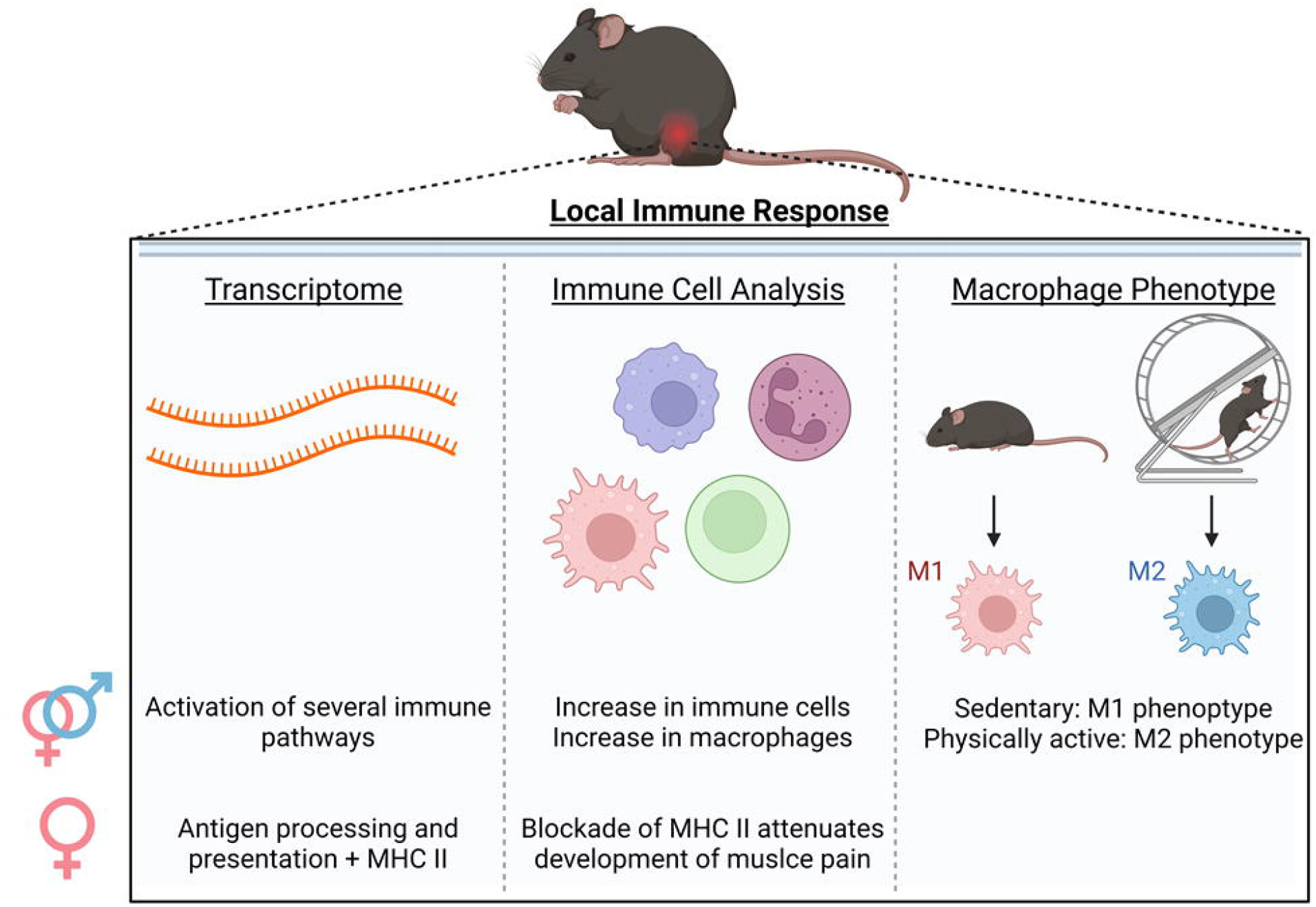

## References

1. D. J. Gaskin, P. Richard, The economic costs of pain in the United States. J Pain 13, 715–724 (2012).

2. M. A. Cimmino, C. Ferrone, M. Cutolo, Epidemiology of chronic musculoskeletal pain. Best Practice & Research Clinical Rheumatology 25, 173–183 (2011).

3. K. J. Berkley, Sex differences in pain. Behav. Brain Sci. 20, 371–380; discussion 435-513 (1997).

4. A. M. Unruh, Gender variations in clinical pain experience. Pain 65, 123–167 (1996).

5. J. S. Mogil, Qualitative sex differences in pain processing: emerging evidence of a biased literature. Nature reviews. Neuroscience 21, 353–365 (2020).

6. A. M. Cowie, B. N. Dittel, C. L. Stucky, A Novel Sex-Dependent Target for the Treatment of Postoperative Pain: The NLRP3 Inflammasome. Front. Neurol. 10, 622–622 (2019).

7. G. Baskozos et al., Comprehensive analysis of long noncoding RNA expression in dorsal root ganglion reveals cell-type specificity and dysregulation after nerve injury. Pain 160, 463–485 (2019).

8. J. Mecklenburg et al., Sex-dependent pain trajectories induced by prolactin require an inflammatory response for pain resolution. Brain. Behav. Immun. 101, 246–263 (2022).

9. R. E. Sorge et al., Different immune cells mediate mechanical pain hypersensitivity in male and female mice. Nat. Neurosci. 18, 1081–1083 (2015).

10. M. Costigan et al., T-cell infiltration and signaling in the adult dorsal spinal cord is a major contributor to neuropathic pain-like hypersensitivity. The Journal of neuroscience : the official journal of the Society for Neuroscience 29, 14415–14422 (2009).

11. W. Y. Gong, R. E. Abdelhamid, C. S. Carvalho, K. A. Sluka, Resident Macrophages in Muscle Contribute to Development of Hyperalgesia in a Mouse Model of Noninflammatory Muscle Pain. J Pain 17, 1081–1094 (2016).

12. N. S. Gregory, R. G. Brito, M. C. Fusaro, K. A. Sluka, ASIC3 Is Required for Development of Fatigue-Induced Hyperalgesia. Molecular neurobiology 53, 1020–1030 (2016).

13. G. de Azambuja et al., Regular swimming exercise prevented the acute and persistent mechanical muscle hyperalgesia by modulation of macrophages phenotypes and inflammatory cytokines via PPARγ receptors. Brain, behavior, and immunity 95, 462–476 (2021).

14. M. C. Oliveira-Fusaro et al., P2X4 Receptors on Muscle Macrophages Are Required for Development of Hyperalgesia in an Animal Model of Activity-Induced Muscle Pain. Molecular neurobiology 57, 1917–1929 (2020).

15. F. Bobinski, J. M. Teixeira, K. A. Sluka, A. R. S. Santos, Interleukin-4 mediates the analgesia produced by low-intensity exercise in mice with neuropathic pain. Pain 159, 437–450 (2018).

16. Y. Tu, M. M. Muley, S. Beggs, M. W. Salter, Microglia-independent peripheral neuropathic pain in male and female mice. Pain 163, (2022).

17. A. Leung, N. S. Gregory, L. A. Allen, K. A. Sluka, Regular physical activity prevents chronic pain by altering resident muscle macrophage phenotype and increasing interleukin-10 in mice. Pain 157, 70–79 (2016).

18. R. Sabharwal, L. Rasmussen, K. A. Sluka, M. W. Chapleau, Exercise prevents development of autonomic dysregulation and hyperalgesia in a mouse model of chronic muscle pain. Pain 157, 387–398 (2016).

19. L. V. Lima, J. M. DeSantana, L. A. Rasmussen, K. A. Sluka, Short-duration physical activity prevents the development of activity-induced hyperalgesia through opioid and serotoninergic mechanisms. Pain 158, 1697–1710 (2017).

20. K. A. Sluka, J. M. O’Donnell, J. Danielson, L. A. Rasmussen, Regular physical activity prevents development of chronic pain and activation of central neurons. J Appl Physiol (1985) 114, 725–733 (2013).

21. C. A. Staunton et al., Skeletal muscle transcriptomics identifies common pathways in nerve crush injury and ageing. Skeletal Muscle 12, 3 (2022).

22. S. L. Asche-Godin et al., RNA-sequencing Reveals a Gene Expression Signature in Skeletal Muscle of a Mouse Model of Age-associated Postoperative Functional Decline. The Journals of Gerontology: Series A 77, 1939–1950 (2022).

23. D. W. McKellar et al., Large-scale integration of single-cell transcriptomic data captures transitional progenitor states in mouse skeletal muscle regeneration. Communications Biology 4, 1280 (2021).

24. N. S. Gregory, K. Gibson-Corley, L. Frey-Law, K. A. Sluka, Fatigue-enhanced hyperalgesia in response to muscle insult: induction and development occur in a sex-dependent manner. Pain 154, 2668–2676 (2013).

25. J. B. Lesnak, S. Inoue, L. Lima, L. Rasmussen, K. A. Sluka, Testosterone protects against the development of widespread muscle pain in mice. Pain 161, 2898–2908 (2020).

26. J. B. Lesnak et al., Resistance training protects against muscle pain through activation of androgen receptors in male and female mice. Pain 163, 1879–1891 (2022).

27. R. V. Contarteze, B. Manchado Fde, C. A. Gobatto, M. A. De Mello, Stress biomarkers in rats submitted to swimming and treadmill running exercises. Comparative biochemistry and physiology. Part A, Molecular & integrative physiology 151, 415–422 (2008).

28. R. G. Brito, L. A. Rasmussen, K. A. Sluka, Regular physical activity prevents development of chronic muscle pain through modulation of supraspinal opioid and serotonergic mechanisms. Pain Rep 2, e618 (2017).

29. Y. Luo et al., New developments on the Encyclopedia of DNA Elements (ENCODE) data portal. Nucleic Acids Res. 48, D882–d889 (2020).

30. An integrated encyclopedia of DNA elements in the human genome. Nature 489, 57–74 (2012).

31. D. Kim, J. M. Paggi, C. Park, C. Bennett, S. L. Salzberg, Graph-based genome alignment and genotyping with HISAT2 and HISAT-genotype. Nat. Biotechnol. 37, 907–915 (2019).

32. D. Kim, B. Langmead, S. L. Salzberg, HISAT: a fast spliced aligner with low memory requirements. Nature Methods 12, 357–360 (2015).

33. R. Patro, G. Duggal, M. I. Love, R. A. Irizarry, C. Kingsford, Salmon provides fast and bias-aware quantification of transcript expression. Nat Methods 14, 417–419 (2017).

34. P. Ewels, M. Magnusson, S. Lundin, M. Käller, MultiQC: summarize analysis results for multiple tools and samples in a single report. Bioinformatics 32, 3047–3048 (2016).

35. C. Soneson, M. I. Love, M. D. Robinson, Differential analyses for RNA-seq: transcript-level estimates improve gene-level inferences. F1000Res 4, 1521 (2015).

36. M. I. Love, W. Huber, S. Anders, Moderated estimation of fold change and dispersion for RNA-seq data with DESeq2. Genome Biol. 15, 550 (2014).

37. M. I. Love, S. Anders, H. Wolfgang. (2022).

38. Y. Benjamini, Y. Hochberg, Controlling the False Discovery Rate: A Practical and Powerful Approach to Multiple Testing. Journal of the Royal Statistical Society. Series B (Methodological) 57, 289–300 (1995).

39. P. Khatri, S. Drăghici, Ontological analysis of gene expression data: current tools, limitations, and open problems. Bioinformatics 21, 3587–3595 (2005).

40. S. Drăghici et al., A systems biology approach for pathway level analysis. Genome Res. 17, 1537–1545 (2007).

41. T.-M. Nguyen, A. Shafi, T. Nguyen, S. Drăghici, Identifying significantly impacted pathways: a comprehensive review and assessment. Genome Biol. 20, 203 (2019).

42. D. A. Skyba, R. Radhakrishnan, K. A. Sluka, Characterization of a method for measuring primary hyperalgesia of deep somatic tissue. J Pain 6, 41–47 (2005).

43. K. A. Sluka, L. A. Rasmussen, Fatiguing exercise enhances hyperalgesia to muscle inflammation. Pain 148, 188–197 (2010).

44. V. Steimle, C. A. Siegrist, A. Mottet, B. Lisowska-Grospierre, B. Mach, Regulation of MHC class II expression by interferon-gamma mediated by the transactivator gene CIITA. Science 265, 106–109 (1994).

45. N. Pishesha, T. J. Harmand, H. L. Ploegh, A guide to antigen processing and presentation. Nature Reviews Immunology, (2022).

46. M. J. Lacagnina, L. R. Watkins, P. M. Grace, Toll-like receptors and their role in persistent pain. Pharmacol. Ther. 184, 145–158 (2018).

47. A. M. Cowie, A. D. Menzel, C. O’Hara, M. W. Lawlor, C. L. Stucky, NOD-like receptor protein 3 inflammasome drives postoperative mechanical pain in a sex-dependent manner. Pain 160, 1794–1816 (2019).

48. L. Leung, C. M. Cahill, TNF-alpha and neuropathic pain--a review. J. Neuroinflammation 7, 27 (2010).

49. F. A. White, N. M. Wilson, Chemokines as pain mediators and modulators. Curr. Opin. Anaesthesiol. 21, 580–585 (2008).

50. D. C. Fritzinger, D. E. Benjamin, The Complement System in Neuropathic and Postoperative Pain. Open Pain J. 9, 26–37 (2016).

51. Y. Kimura et al., IL-33 induces orofacial neuropathic pain through Fyn-dependent phosphorylation of GluN2B in the trigeminal spinal subnucleus caudalis. Brain. Behav. Immun. 99, 266–280 (2022).

52. Z. Song et al., STAT1 as a downstream mediator of ERK signaling contributes to bone cancer pain by regulating MHC II expression in spinal microglia. Brain. Behav. Immun. 60, 161–173 (2017).

53. M. Hartlehnert et al., Schwann cells promote post-traumatic nerve inflammation and neuropathic pain through MHC class II. Sci. Rep. 7, 12518 (2017).

54. H. Liu et al., Immunodominant fragments of myelin basic protein initiate T cell-dependent pain. J. Neuroinflammation 9, 119 (2012).

55. S. Noor et al., Prenatal alcohol exposure potentiates chronic neuropathic pain, spinal glial and immune cell activation and alters sciatic nerve and DRG cytokine levels. Brain, behavior, and immunity 61, 80–95 (2017).

56. S. M. Sweitzer, K. A. White, C. Dutta, J. A. DeLeo, The differential role of spinal MHC class II and cellular adhesion molecules in peripheral inflammatory versus neuropathic pain in rodents. J. Neuroimmunol. 125, 82–93 (2002).

57. H. Hashizume, M. D. Rutkowski, J. N. Weinstein, J. A. DeLeo, Central administration of methotrexate reduces mechanical allodynia in an animal model of radiculopathy/sciatica. Pain 87, 159–169 (2000).

58. S. M. Sweitzer, J. A. DeLeo, The active metabolite of leflunomide, an immunosuppressive agent, reduces mechanical sensitivity in a rat mononeuropathy model. The Journal of Pain 3, 360–368 (2002).

59. A. Moss et al., Spinal microglia and neuropathic pain in young rats. Pain 128, (2007).

60. P. Hu, A. L. Bembrick, K. A. Keay, E. M. McLachlan, Immune cell involvement in dorsal root ganglia and spinal cord after chronic constriction or transection of the rat sciatic nerve. Brain. Behav. Immun. 21, 599–616 (2007).

61. Z. Li et al., Spinal versus brain microglial and macrophage activation traits determine the differential neuroinflammatory responses and analgesic effect of minocycline in chronic neuropathic pain. Brain, behavior, and immunity 58, 107–117 (2016).

62. P. Cruz-Tapias, J. Castiblanco, J. Anaya, Major histocompatibility complex: Antigen processing and presentation. Autoimmunity: From Bench to Bedside (El Rosario University Press, 2013).

63. M. Wieczorek et al., Major Histocompatibility Complex (MHC) Class I and MHC Class II Proteins: Conformational Plasticity in Antigen Presentation. Front. Immunol. 8, (2017).

64. E. H. Jin et al., Genome-wide expression profiling of complex regional pain syndrome. PLoS One 8, e79435 (2013).

65. J. J. van Hilten, W. J. van de Beek, B. O. Roep, Multifocal or generalized tonic dystonia of complex regional pain syndrome: a distinct clinical entity associated with HLA-DR13. Ann. Neurol. 48, 113–116 (2000).

66. M. A. Kemler et al., HLA-DQ1 associated with reflex sympathetic dystrophy. Neurology 53, 1350–1351 (1999).

67. B. Klimenta, H. Nefic, N. Prodanovic, R. Jadric, F. Hukic, Association of biomarkers of inflammation and HLA-DRB1 gene locus with risk of developing rheumatoid arthritis in females. Rheumatology international 39, 2147–2157 (2019).

68. G. M. Cavestro et al., Association of HLA-DRB1*0401 allele with chronic pancreatitis. Pancreas 26, 388–391 (2003).

69. J. Satsangi et al., Contribution of genes of the major histocompatibility complex to susceptibility and disease phenotype in inflammatory bowel disease. Lancet 347, 1212–1217 (1996).

70. P. R. Ray et al., RNA profiling of human dorsal root ganglia reveals sex-differences in mechanisms promoting neuropathic pain. Brain, (2022).

71. S. G. Dorsey et al., Whole blood transcriptomic profiles can differentiate vulnerability to chronic low back pain. PLoS One 14, e0216539 (2019).

72. M. Parisien et al., Effect of Human Genetic Variability on Gene Expression in Dorsal Root Ganglia and Association with Pain Phenotypes. Cell Rep. 19, 1940–1952 (2017).

73. Z. Chen et al., Regulatory T cells-centered regulatory networks of skeletal muscle inflammation and regeneration. Cell Biosci. 12, 112 (2022).

74. J. Zhang et al., CD8 T cells are involved in skeletal muscle regeneration through facilitating MCP-1 secretion and Gr1(high) macrophage infiltration. J. Immunol. 193, 5149–5160 (2014).

75. D. Burzyn et al., A special population of regulatory T cells potentiates muscle repair. Cell 155, 1282–1295 (2013).

76. A. Castiglioni et al., FOXP3+ T Cells Recruited to Sites of Sterile Skeletal Muscle Injury Regulate the Fate of Satellite Cells and Guide Effective Tissue Regeneration. PLoS One 10, e0128094 (2015).

77. G. Moalem, K. Xu, L. Yu, T lymphocytes play a role in neuropathic pain following peripheral nerve injury in rats. Neuroscience 129, 767–777 (2004).

78. E. J. Cobos et al., Mechanistic Differences in Neuropathic Pain Modalities Revealed by Correlating Behavior with Global Expression Profiling. Cell Rep. 22, 1301–1312 (2018).

79. P. M. Grace et al., Prior voluntary wheel running attenuates neuropathic pain. Pain 157, 2012–2023 (2016).

80. L. R. Watkins et al., Targeted interleukin-10 plasmid DNA therapy in the treatment of osteoarthritis: Toxicology and pain efficacy assessments. Brain, behavior, and immunity 90, 155–166 (2020).

81. P. M. Grace et al., Behavioral assessment of neuropathic pain, fatigue, and anxiety in experimental autoimmune encephalomyelitis (EAE) and attenuation by interleukin-10 gene therapy. Brain. Behav. Immun. 59, 49–54 (2017).

82. A. Ledeboer et al., Intrathecal interleukin-10 gene therapy attenuates paclitaxel-induced mechanical allodynia and proinflammatory cytokine expression in dorsal root ganglia in rats. Brain. Behav. Immun. 21, 686–698 (2007).

83. E. Sloane et al., Anti-inflammatory cytokine gene therapy decreases sensory and motor dysfunction in experimental Multiple Sclerosis: MOG-EAE behavioral and anatomical symptom treatment with cytokine gene therapy. Brain. Behav. Immun. 23, 92–100 (2009).

84. F. Bobinski et al., Role of brainstem serotonin in analgesia produced by low-intensity exercise on neuropathic pain after sciatic nerve injury in mice. Pain 156, 2595–2606 (2015).

85. S. K. Mittal, P. A. Roche, Suppression of antigen presentation by IL-10. Curr. Opin. Immunol. 34, 22–27 (2015).

86. S. Sorichter et al., Skeletal troponin I as a marker of exercise-induced muscle damage. J Appl Physiol (1985) 83, 1076–1082 (1997).

87. S. M. Green-Fulgham et al., Preconditioning by voluntary wheel running attenuates later neuropathic pain via nuclear factor E2–related factor 2 antioxidant signaling in rats. Pain 163, (2022).

88. M. Abdullah et al., Gender effect on in vitro lymphocyte subset levels of healthy individuals. Cell. Immunol. 272, 214–219 (2012).

89. S. Sankaran-Walters et al., Sex differences matter in the gut: effect on mucosal immune activation and inflammation. Biol. Sex Differ. 4, 10 (2013).

90. A. C. Roden et al., Augmentation of T cell levels and responses induced by androgen deprivation. J. Immunol. 173, 6098–6108 (2004).

91. A. A. Lin, S. E. Wojciechowski, D. A. Hildeman, Androgens suppress antigen-specific T cell responses and IFN-γ production during intracranial LCMV infection. J. Neuroimmunol. 226, 8–19 (2010).

92. H. Lotter et al., Testosterone increases susceptibility to amebic liver abscess in mice and mediates inhibition of IFNγ secretion in natural killer T cells. PLoS One 8, e55694 (2013).

93. S. L. Klein, K. L. Flanagan, Sex differences in immune responses. Nature Reviews Immunology 16, 626–638 (2016).

94. R. E. Sorge, S. K. Totsch, Sex Differences in Pain. J. Neurosci. Res. 95, 1271–1281 (2017).

