## Supplementary figures and images for "The impact of sex and physical activity on the local immune response to muscle pain"

### Supplemental Figure 1

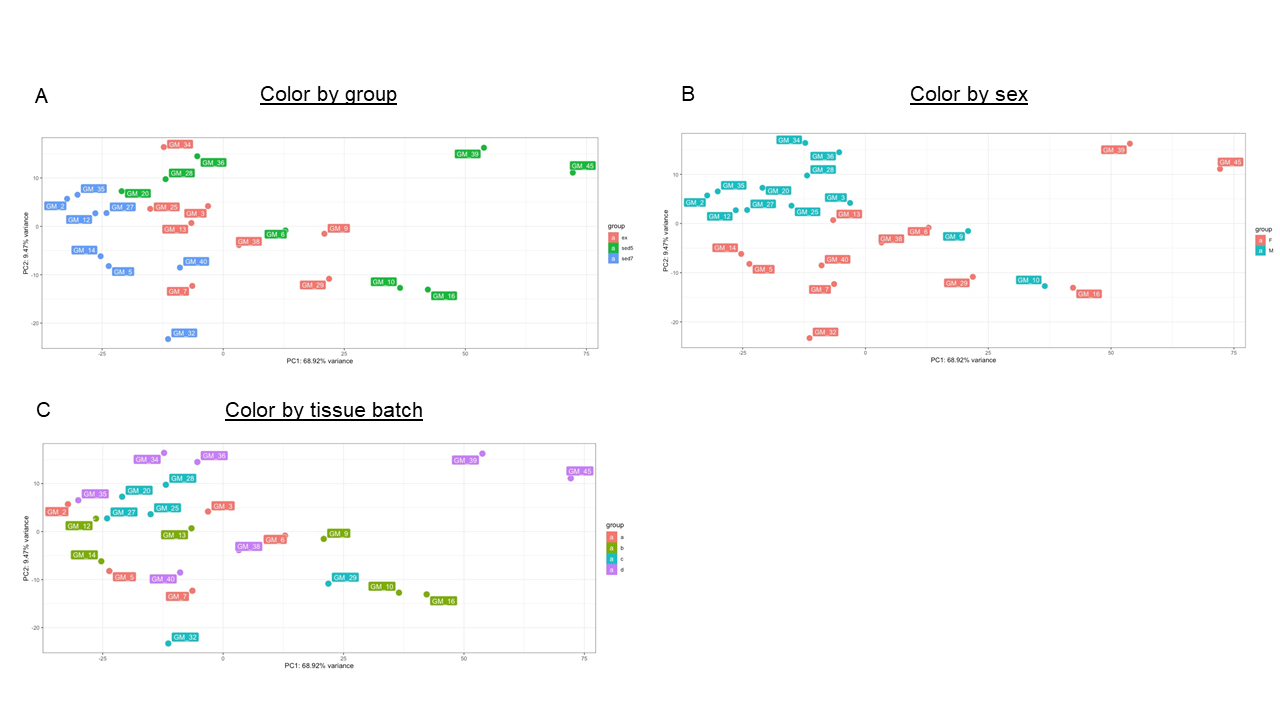

### Supplemental Figure 2

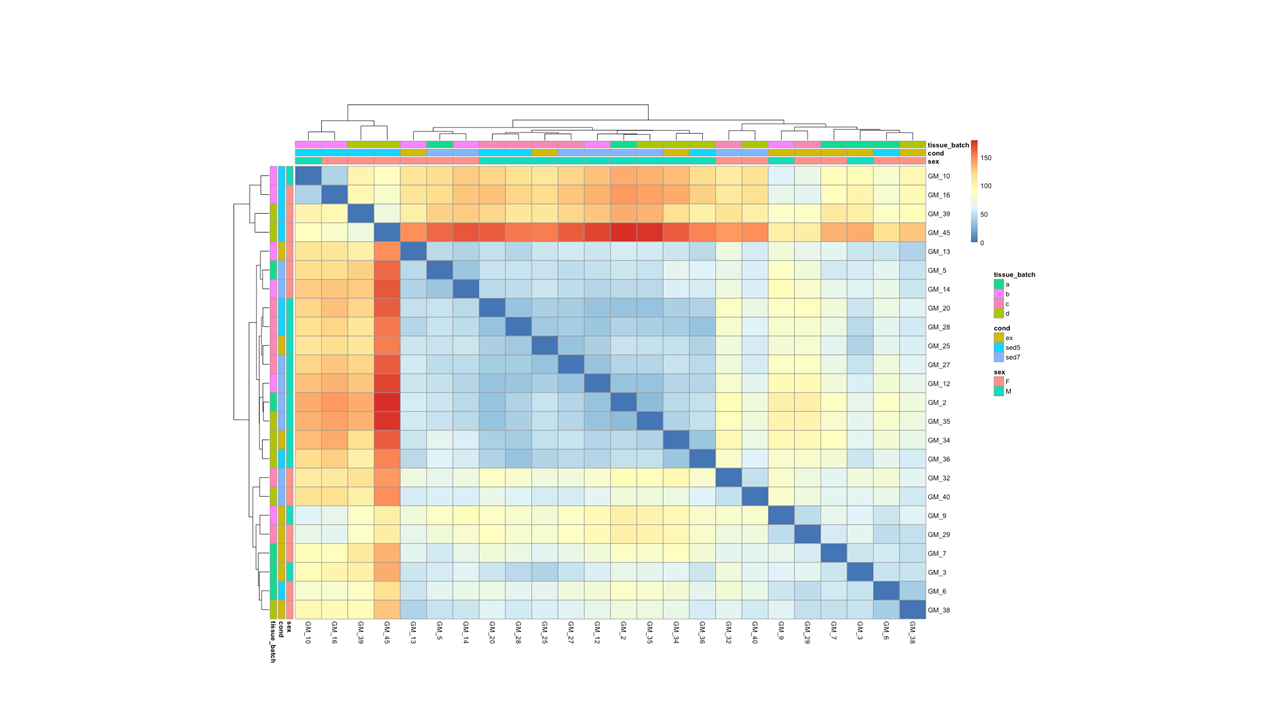

### Supplemental Table 1

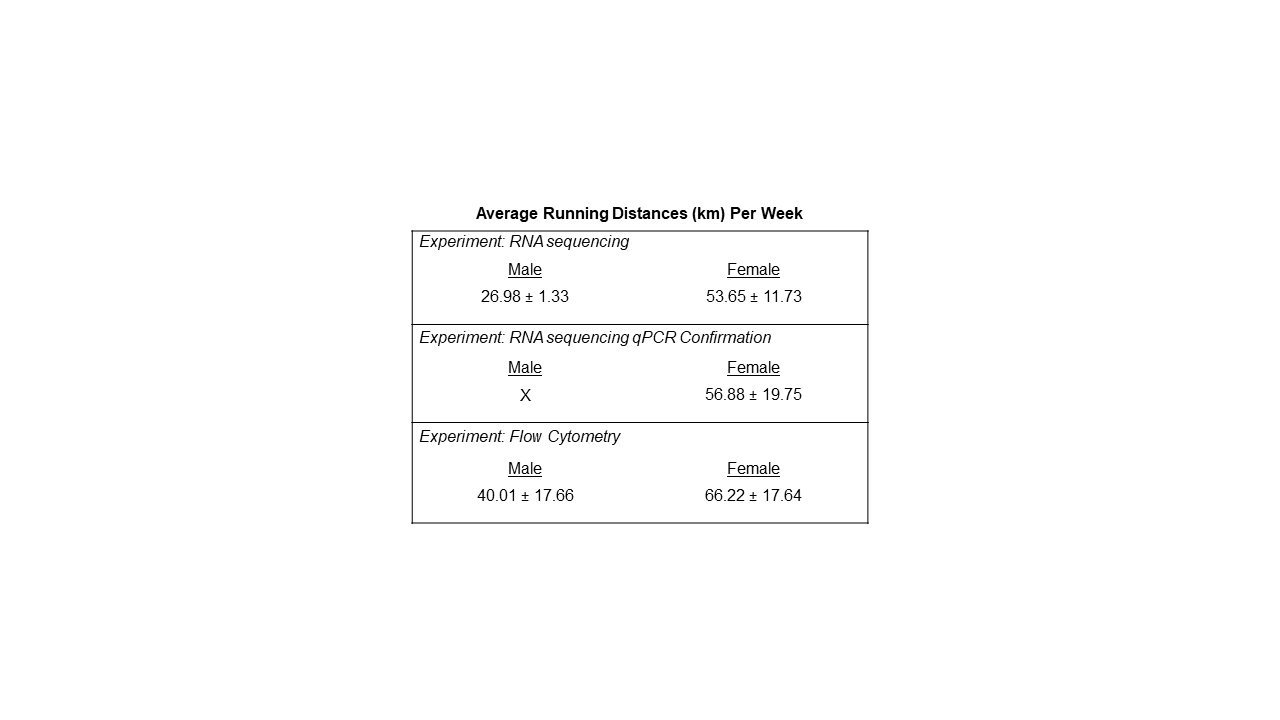

### Supplemental Table 2

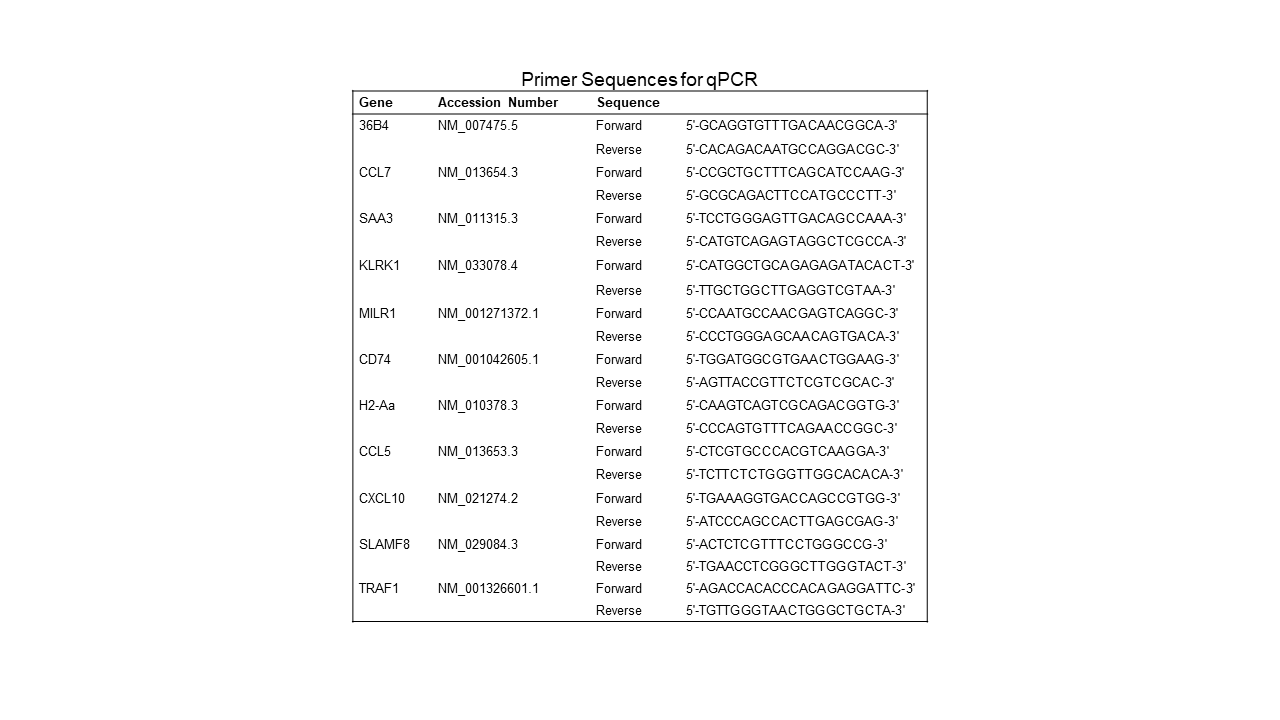

### Supplemental Table 3

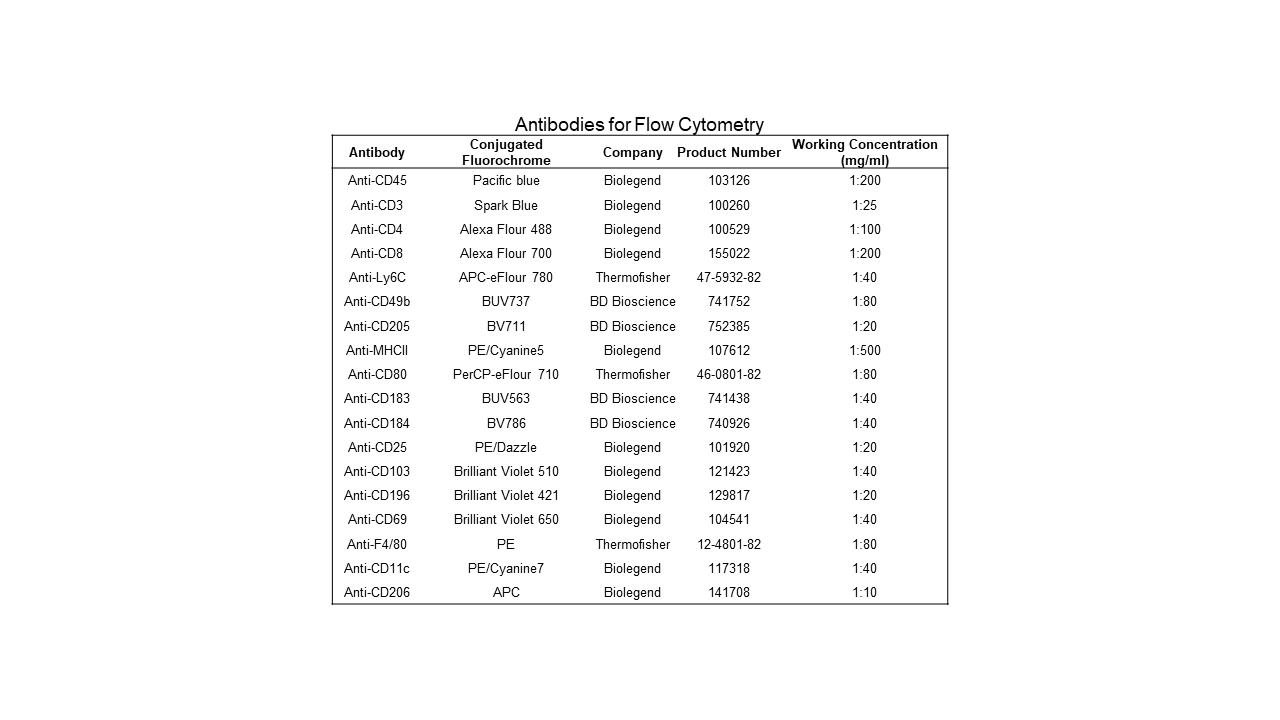
